# Oxytocin-like signaling couples reproductive state to intestinal lipid metabolism in aging *C. elegans*

**DOI:** 10.64898/2026.06.28.735098

**Authors:** Katelyn M. Adam, Kathryn M. Kuklinski, Charles A. Fisher, Will Skinner, Jacqueline Y. Lo, Alison Kochersberger, Jennifer L. Garrison

## Abstract

Oxytocin and vasopressin are endogenous bioactive peptides with conserved roles in reproduction and, more recently recognized, in peripheral lipid metabolism. Whether this signaling system also shapes how reproduction declines with age has not been tested in any animal. Here we show that in *C. elegans*, the oxytocin/vasopressin-like neuropeptide nematocin restrains reproductive output as animals reach mid-life. Nematocin and its two receptors are produced throughout adult life and peak as reproduction begins to wane. Animals lacking receptor signaling produce more offspring in mid-life, an improvement that reflects better egg quality and fertilization rather than improved embryo survival. This benefit is accompanied by changes in intestinal fat metabolism, the worm’s equivalent of liver and adipose tissue: nematocin normally limits the activity of a fatty-acid desaturase that is otherwise induced by mating, and it shapes how much yolk reaches developing eggs. The two receptors act through separate routes, one tuning intestinal fat metabolism and the other controlling yolk delivery to the egg. Together, these findings reveal nematocin as a regulator of the intestinal metabolic environment across reproductive age, mirroring the recently described oxytocin-hepatocyte-adipocyte lipid axis in mammals and implicate this conserved signaling system in the coordination of maternal investment during reproductive aging.

## Introduction

Reproductive lifespan depends on the availability of sex cells (gametes). The quantity, quality, and temporal distribution of gametes define reproductive strategy across animals and are coordinated by both local gonadal signals and systemic physiological regulation. Gamete production is governed largely within the gonad. However, reproductive timing and investment must track environmental conditions, nutritional state, and somatic health. Such coordination demands extensive inter-tissue communication and higher-level neuroendocrine control.

Reproductive aging is characterized by an age-dependent decline in oocyte quality and number^1–9^. In many species reproductive decline is accompanied by altered neuroendocrine signaling^10,11,11–17^, partially precipitated by diminishing gonadal feedback. However, permissive factors can modify the onset and severity of germline deterioration^1,5,18–32^, suggesting that the reproductive decline is plastic and receptive to systemic regulation. This continued reliance on external signals affords physiological flexibility but also introduces vulnerability. Determining which neuroendocrine pathways remain functional, become sensitized, or become deregulated with reproductive age is therefore an important and underexplored question in reproductive aging biology.

Reproduction is energetically expensive, and reproductive aging may reflect an age-dependent failure to allocate somatic resources. Oocyte production, fertilization, embryonic provisioning, and lactation are metabolically costly processes that rely heavily on lipid mobilization and transport^20,33–37^. Maternal provisioning is fundamentally a problem of nutrient allocation, and successful reproduction depends on the ability of peripheral metabolic tissues to synthesize, store, and redistribute lipids in response to reproductive demand. Metabolic dysfunction is associated with reproductive decline, subfertility, and reproductive pathologies^38–62^. Yet the directionality of this relationship remains unresolved: metabolic dysfunction may contribute to reproductive decline, reproductive aging may disrupt metabolic homeostasis, or shared neuroendocrine signals may control both. Understanding how metabolism is regulated across the reproductive span is therefore essential to understanding reproductive aging itself.

Among the neuroendocrine systems capable of coordinating reproduction and metabolism, the oxytocin/vasopressin (OT/VP) peptide family is uniquely positioned as a candidate regulator of reproductive aging. Oxytocin and vasopressin are ancient neuropeptides conserved across bilaterians and best known for their roles in parturition^63–66^, lactation^65,67,68^, social bonding^69,70^, reproductive^71^ and feeding behaviors^72–74^. However, their functions extend beyond these classical roles. OT/VP signaling also regulates ovulation timing^49,75–81^, steroid biosynthesis^82,83^, sperm physiology^84,85^, embryonic diapause^86^, and reproductive tract function^87,88^. Less appreciated are their roles in peripheral metabolism, including the regulation of adipose lipolysis^89,90^, hepatic lipid metabolism^46,91–93^, insulin sensitivity^94–96^, bone mineralization^97,98^ and intestinal physiology^99–104^. This dual role in both reproductive physiology and systemic metabolism make OT/VP signaling particularly compelling as a mechanism through which reproductive state could influence maternal metabolic investment with age.

The evolutionary conservation of this signaling system allows these questions to be addressed in a genetically tractable model with a stereotyped reproductive span and well-defined mechanisms of reproductive aging^6^. In *C. elegans*, nematocin, *ntc-1,* and its receptors, *ntr-1* and *ntr-2,* constitute an oxytocin/vasopressin-like neuropeptide signaling system. Nematocin is known to regulate male mating behavior^105^, social feeding^106^, and associative gustatory learning^107^, but its role in maternal provisioning, intestinal lipid metabolism, and reproductive aging remains unknown. Importantly, nematocin signaling is positioned within a broader neuroendocrine network with access to peripheral metabolic tissues^108–111^, creating the potential for neuroendocrine regulation of reproductive investment through control of somatic metabolic pathways.

*Caenorhabditis elegans* offers a tractable system for distinguishing these possibilities, because mating drives a rapid and quantifiable redistribution of maternal resources toward reproduction^112–115^. Mating triggers physiological changes that increase reproductive success at the expense of somatic maintenance and maternal lifespan^112,113,116–119^ . Male-derived pheromones increase germline activity and improve oocyte quality^120–124^, while seminal fluid proteins promote lipid mobilization into yolk production and bias lipid desaturation in ways that facilitate sperm migration^115^. Sperm-associated proteins stimulate ovulation^125–128^ and increase fertilizability^129,130^ through signaling pathways that enhance reproductive output^131–133^. Together, these responses constitute a coordinated post-mating response that actively prioritizes reproductive investment over maternal somatic maintenance.

Over the course of the reproductive span, the metabolic demand imposed by yolk production progressively depletes intestinal lipid stores and contributes to intestinal atrophy and maternal decline^134–137^. This depletion is driven in large part by the requirement for polyunsaturated fatty acids (PUFAs)^115^, which are essential for oocyte production, membrane composition, sperm guidance^138,139^, and nutritional provisioning to post-embryonic development and progeny resilence^135,140–144^. Lipid desaturation therefore represents more than a metabolic consequence of reproduction; it is a central determinant of reproductive success and maternal investment. Importantly, mating-induced changes in lipid metabolism are not simply passive responses to oocyte production. Lipid desaturases are differentially regulated after mating, and the stearoyl-CoA desaturase FAT-7, the functional homolog of mammalian SCD1, is transcriptionally upregulated even in animals defective in oogenesis^112^.Thus the lipid demand imposed by oocyte production is uncoupled from the transcriptional reprogramming that is specific to mating-induced lipid remodeling.

FAT-7 catalyzes the production of oleic acid, a monounsaturated fatty acid subsequently shown to be important for maintaining maternal lipid reserves after mating^112^. Because oleic acid supports both reproductive metabolism and maternal somatic maintenance, FAT-7 abundance serves as a meaningful proxy for the intestinal metabolic state relevant to the mated response and maternal provisioning. In *C. elegans*, the intestine performs the metabolic functions of both the mammalian liver and adipose tissue, including lipid synthesis, storage, and secretion^145^. It is also the site of vitellogenin production^146^, the process through which maternal nutrients are packaged for oocyte uptake.

Vitellogenins, functionally analogous to mammalian lipoproteins such as low-density lipoproteins, are synthesized in the intestine, secreted into the body cavity, and taken up by developing oocytes through receptor-mediated endocytosis^147^. As a result, intestinal lipid metabolism sits at the center of reproductive investment, linking somatic metabolic state directly to fertility and maternal provisioning.

Preliminary evidence further supports this possibility. Published microarray data show that *ntr-1* and *ntr-2* expression is downregulated by mating in young adults^112^, suggesting that reproductive state alters receptor abundance and potentially nematocin sensitivity. This observation raises the possibility that mating and reproductive age dynamically shape nematocin signaling, but whether this relationship influences intestinal metabolism or fertility has not been characterized.

We hypothesized that nematocin signaling balances maternal lipid metabolism with reproductive investment as animals age. Specifically, we asked whether the age-associated rise in nematocin expression is associated with changes in intestinal lipid desaturation, maternal yolk provisioning, and mated fertility. Using loss-of-function alleles of *ntc-1, ntr-1*, and *ntr-2*, individually and in combination, we measured nematocin pathway action across adulthood, mated fertility, intestinal lipid desaturation, and yolk abundance during peak reproduction and reproductive decline.

Our results reveal that loss of nematocin signaling improves mated fecundity during reproductive decline in an *ntr-1*-dependent manner, elevates intestinal FAT-7 abundance, and alters yolk provisioning in a temporally dynamic, age-dependent fashion. Together, these data suggest nematocin acts as a neuroendocrine brake on maternal metabolic investment, balancing reproductive output and nutritional provisioning against the intestinal metabolic environment with age. These findings parallel the recently described oxytocin–hepatocyte-adipocyte lipid axis in mammals^89,91,148^ and establish a foundation for testing how neuroendocrine control of lipid metabolism shapes fertility decline with age. More broadly, this work supports the idea that reproductive aging is not solely a consequence of germline depletion, but also a systems-level process shaped by conserved neuroendocrine regulation of somatic metabolism.

## Results

### Nematocin signaling genes are dynamically expressed across reproductive life stages and their loss is associated with increased mated fecundity during reproductive decline

#### Nematocin signaling is highest at the onset of reproductive decline

Although the oxytocin/vasopressin signaling family is ancient and broadly conserved, its temporal relationship to reproductive aging has not been systematically characterized in any model organism. To determine whether nematocin signaling changes across the reproductive span, we measured *ntc-1*, *ntr-1*, and *ntr-2* transcript abundance across adulthood in age-synchronized wild-type animals and expressed each relative to day 1 of adulthood (**Figure 1A**).

**Figure 1:**
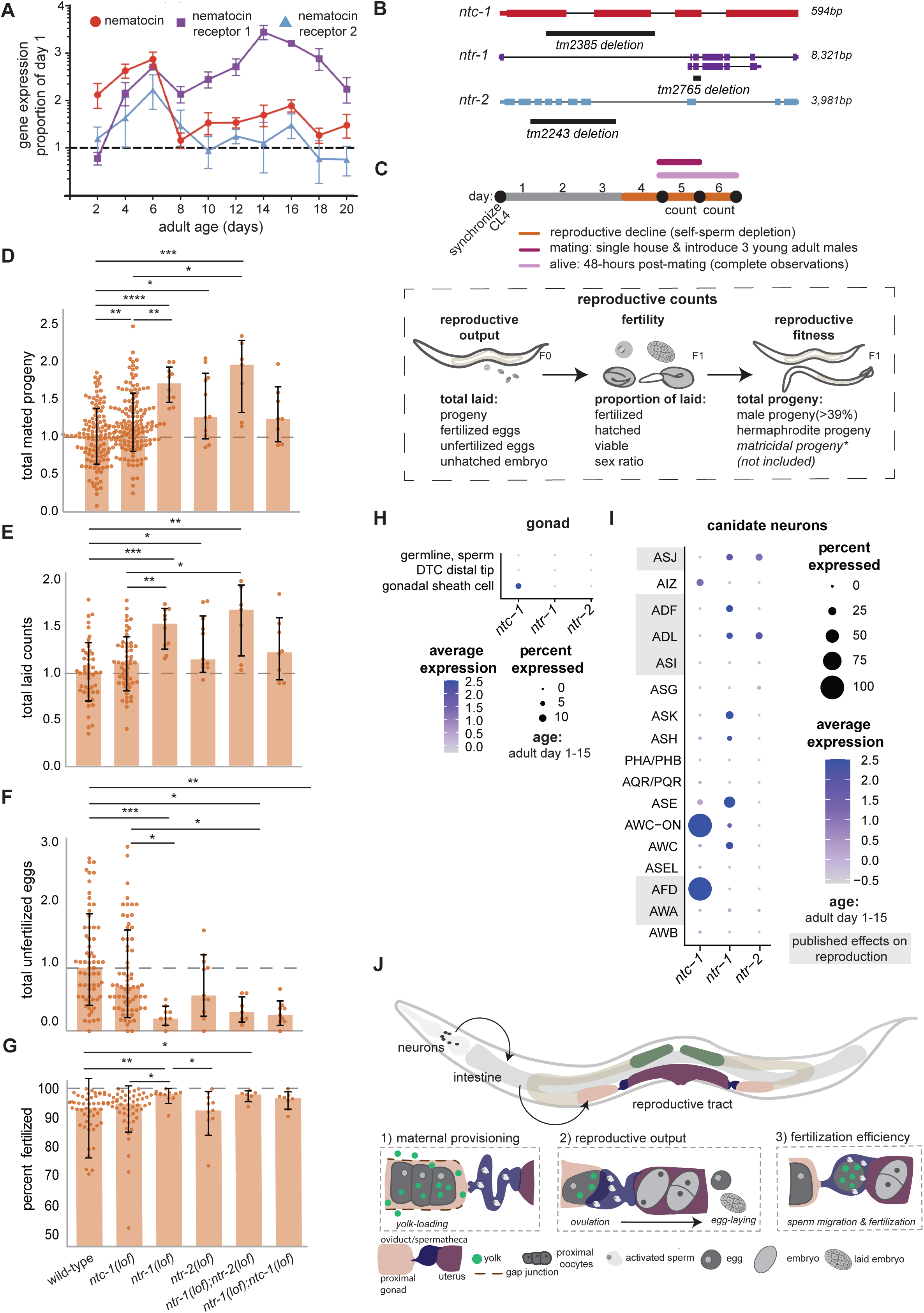
Elevated nematocin signaling is associated with the age-related decline in reproductive capacity. **(A)** Gene expression of *ntc-1*, *ntr-1* and *ntr-2* measured by qPCR throughout adulthood of wild-type N2 animals. Relative expression normalized to the first day of adulthood for each gene. (N = 3, n = > 100 worms) **(B)** Schematic of *ntc-1*, *ntr-1* and *ntr-2* gene structure, exons (boxes) and introns (lines), with annotations of alleles used in this study (black bars). All deletion alleles are loss of function (lof). **(C)** Experimental design of mated fertility during the reproductive decline(D-G). Animals were synchronized by picking crescent L4s, aged in groups of 50 or singly housed until mated with 3 young adult males at the start of adult day 5, shown as a dark red line. Mating occurred for 24 hours and animals were transferred to new plates every 24 hours to count reproductive output. Mated worms were only included if they survived for 48 hours from the start of the assay. Metrics evaluated are shown below. **(D)** Relative total mated progeny counts from adult days 5 and 6. **(E)** Relative total mated laid counts from adult days 5-6. Laid counts are the sum of all progeny, unfertilized and unhatched eggs during the collection interval. **(F)** Relative total mated unfertilized eggs from adult days 5-6. **(G)** Proportion of fertilized eggs after mating from adult days 5-6. **(H)** Single-cell dot plot of gonadal cell types and **(I)** candidate neurons normalized to the relative expression of *ntc-1*, *ntr-1* and *ntr-2* across adult days 1-15. Data obtained from Calico aging sc-RNA-seq dataset.^1^ **(I)** Schematic of proposed model for increased mid-life reproductive capacity. We hypothesized that nematocin signaling, likely from neurons, remodels intestinal lipid metabolism, influencing maternal provisioning and consequently reproductive capacity with age. Depicted are three reproductive mechanisms changed by loss of nematocin signaling, which directly impact fertility and post-embryonic progeny resilience. Maternal resources are provisioned from the intestine and delivered to the oocyte in the form of yolk (Box 1). Yolk (green) is rich in lipid substrates important for sperm guidance and subsequently fertilization, a determinant of oocyte quality shown in Box 3. After each ovulation, sperm are pushed out of the oviduct/spermatheca and must follow oocyte-derived cues to migrate back to successfully fertilize the next oocyte. Insufficient signaling by the oocyte results in sperm mis-localization and decreased fertilization efficiency. Sperm secrete major sperm proteins (MSPs) that promote oocyte maturation, gonad contraction and oocyte entry into the oviduct/spermatheca. In their absence, oogenesis is stalled. Reproductive output is dependent upon steady oocyte production, ovulation and egg-laying (Box 2). Changes to reproductive output when egg-laying is normal are primarily caused by mechanisms controlling oogenesis. **(D-G)** All conditions were normalized to the median wild-type control. Kruskal- Wallis + Dunn Test + Benjamini-Hochberg (FDR) corrections were applied to pooled samples between all genotypes. All significant comparisons and adjusted p-values are shown. Adjusted *p*-values indicated by * *p* < 0.05, ** *p* <0.01, *** *p* < 0.001, **** *p* < 0.0001. Number of experimental (N) and biological replicates(n) shown **(D)** N = 8, 8, 1, 2,1, 1; n = 90, 105, 10, 11, 7, 8 **(E)** N = 3, 3, 1, 2, 1, 1; n = 17, 20,10,11,7,8 **(F)** N = 4,4,1,2,1,1; n = 33, 33, 9, 11, 7,9 **(G)** N = 3, 3, 1, 2, 1, 1; n = 17, 20, 10, 11, 7, 8

All three genes were elevated above the day 1 baseline throughout the reproductive period, consistent with the established roles of oxytocin/vasopressin-like signaling in reproductive physiology. However, expression did not peak during the period of maximal oocyte production. Instead, all three genes reached maximal abundance during the onset of reproductive decline, on adult days 4–6. This temporal pattern suggests that nematocin signaling is most active not during peak fertility, but during the transition into declining reproductive capacity.

Following reproductive cessation, receptor expression diverged. *ntr-1* remained elevated through the end of the wild-type lifespan, whereas *ntr-2* and *ntc-1* declined after reproduction had ceased. This receptor-specific divergence is consistent with *ntr-1* becoming the functionally dominant receptor with advancing age, a pattern that recurs throughout our phenotypic analyses.

As in mammals, where neonatal oxytocin antagonism or receptor loss do not broadly compromise early reproductive capacity^149–151^, loss of nematocin signaling in young *C. elegans* does not impair fertility^105,106^ (**Supplementary Figure 1A-I**). The peak of nematocin expression at the onset of reproductive decline, rather than during maximal oocyte production, raises the possibility that it is not required to drive reproduction, but may instead act to limit reproductive capacity during mid-life. This distinction has direct implications for interpreting the loss-of-function phenotypes described below.

#### Loss of *ntr-1* dependent signaling improves mated fecundity during reproductive decline

To test whether elevated nematocin signaling functionally suppresses fertility during reproductive aging, we mated reproductively middle-aged hermaphrodites and quantified viable progeny produced on adult days 5–6 across wild-type animals and a panel of loss-of-function mutants affecting *ntc-1*, *ntr-1*, and *ntr-2* (**Figure 1B**) individually and in combination (**Figure 1C-G & Supplementary Figure 2 A-D**).

Loss of nematocin signaling improved mid-life reproductive output in all genetic backgrounds examined, although the magnitude of the effect depended strongly on receptor genotype. Loss of *ntr-1* alone increased viable progeny by approximately 70% relative to wild type (1 vs 1.7, p < 0.0001; **Figure 1D**), while simultaneous loss of both receptors, *ntr-1;ntr-2*, produced the largest increase, nearly doubling reproductive output (1 vs 1.95, p = 0.0006; **Figure 1D**). By contrast, loss of the peptide itself, *ntc-1,* produced only a modest increase (1 vs 1.16, p = 0.0039; **Figure 1D**), and *ntr-2(lof)* animals showed an intermediate response (1 vs 1.25, p = 0.018; **Figure 1D**). Animals lacking *ntc-1* were phenotypically similar to *ntr-2(lof)* and *ntc-1;ntr-1(lof)* animals (1.16 vs 1.25 and 1.23; **Figure 1D**), but significantly lower than either *ntr-1(lof)* alone or the double receptor mutant (1.16 vs 1.7, p = 0.0039 and 1.16 vs 1.95, p = 0.018; **Figure 1D**).

Together, these data indicate that nematocin signaling suppresses reproductive output during mid-life and that this effect occurs primarily through NTR-1, with NTR-2 making a more modest contribution.

#### Improved fertility reflects increased ovulation and fertilization efficiency rather than embryonic competence

To determine which aspect of reproductive physiology explained the fecundity improvement, we separated total reproductive output into four biologically distinct components: total laid events, unfertilized egg counts, fertilization efficiency, and embryonic competence (**Figure 1D-G & Supplementary Figure 2A-D**). These measurements distinguish whether improved fertility arises from increased ovulation, improved oocyte quality and fertilization, or enhanced post-fertilization embryo viability.

Total laid events reflect ovulatory output, while unfertilized eggs indicate failed fertilization and are commonly associated with reduced oocyte quality^152–156^. Fertilization efficiency measures the proportion of laid eggs that are successfully fertilized, and embryonic competence reflects the developmental quality of embryos after fertilization.

*ntr-1(lof)* and *ntr-1;ntr-2(lof)* animals significantly increased the total number of laid events relative to wild-type(1.00 vs 1.52, p = 0.003 and 1.00 vs 1.67, p = 0.005; **Figure 1E**), and *ntc-1(lof)* animals (0.97 vs 1.52, p = 0.003 and 0.97 vs 1.67, p = 0.005; **Figure 1E**) consistent with elevated ovulation rates. In addition to increased output, both genotypes fertilized a greater proportion of their eggs relative to wild-type (90.8% vs 98.3%, p = 0.004 and 90.8% vs 98.1%, p = 0.015; **Figure 1G**) and produced substantially fewer unfertilized eggs (1.00 vs 0.20, p = 0.0006,1 vs 0.3, p = 0.01, 1.00 vs 0.26, p = 0.001; **Figure 1F**). Loss of *ntr-1*, in any genetic background, strongly reduced unfertilized eggs, whereas loss of *ntc-1* or *ntr-2* alone had little effect relative to wild-type(1.00 vs 0.77, and 1.00 vs 0.66; **Figure 1F**). By contrast, neither metric used to evaluate embryonic competence (percent viability and total unhatched eggs) detected a deviation from wild type (**Supplementary Figure 2B-D**). Because only a small fraction of wild-type embryos (∼3%) were defective at this stage, embryonic quality was unlikely to be a major determinant of fertility in *ntr-1(lof)* or *ntr-1(lof);ntr-2(lof)* animals. However, the total number of unhatched embryos was significantly higher in *ntr-1(lof);ntc-1(lof)* animals (1.00 vs 4.00, p = 0.018; **Supplementary Figure 2C**).

These data indicate that the mid-life fecundity advantage reflects two parallel improvements: increased ovulatory output and enhanced fertilization efficiency. Because embryo quality is unchanged, the phenotype arises before fertilization and is most consistent with improved oocyte quality.

#### Nematocin signaling regulates fertility through extra-gonadal metabolic pathways

To identify the mechanistic basis of this improvement in oocyte quality, we next asked whether nematocin signaling acts directly within reproductive tissues. Surprisingly, tissue-specific expression data did not reliably detect nematocin signaling in the gametes, including the oocyte, and sperm. While, the somatic support cells, analogous to mammalian granulosa and theca cells, only minimally express the peptide, *ntc-1* (**Figure 1H**).

Because neither germ cells nor gonadal support cells show significant nematocin signaling, improved fertility is unlikely to arise from direct gonadal regulation. Instead, these findings point toward systemic pathways capable of altering oocyte quality indirectly, particularly through neuroendocrine regulation of metabolic tissues that determine maternal provisioning. This type of indirect regulation has clear precedent in *C. elegans*, where mating-induced oogenesis and oocyte lipid composition are strongly influenced by neuroendocrine responses to male pheromones^120,121,124^ and population-density cues^140,157–159^, supporting the possibility that nematocin influences reproductive output through peripheral metabolism rather than through the gonad itself.

#### Nematocin signaling is positioned to coordinate peripheral metabolism to reproductive status

Unlike mammals, *C. elegans* lacks a clearly defined hypothalamic–pituitary–gonadal axis. Instead, specific neurons regulate germline activity and oocyte quality in response to mating or social cues (ADL, ASI)^121,122^, nutrient availability (ASJ, AWC), temperature (ASJ, AFD)^160,161^, and stress (ASI)^162,163^.

Nematocin receptors are expressed in many of these candidate neurons, including those known to increase germline activity in response to mating (**Figure 1I**), but are also expressed in neurons known to regulate intestinal lipid metabolism and membrane composition (ADL, ADF, ASK, ASI,AWC,ASJ).^164–171^

Published single-cell expression data support the hypothesis that nematocin signaling could coordinate reproductive status with peripheral metabolism (**Figure 1I**). This possibility is strengthened by the observation that *ntr-1* and *ntr-2* are transcriptionally downregulated in response to mating^112^, while mating itself induces intestinal lipid remodeling. Together, these findings are consistent with a model in which elevated nematocin signaling suppresses, rather than promotes, the metabolic response to mating.

Because improved fertility in nematocin mutants was most consistent with improved oocyte quality, and because oocyte quality is influenced by maternal lipid provisioning, we next asked whether nematocin regulates intestinal lipid desaturation, an important determinant of reproductive metabolism.

### Loss of nematocin signaling is associated with altered intestinal FAT-7 abundance and regional distribution across reproductive age

#### Loss of *ntr-1* increases intestinal FAT-7 abundance independently of age

Mating induces extensive remodeling of intestinal lipid metabolism^112,113,115,118^, including increased lipid turnover, altered fatty acid composition, and enhanced lipid transport to the germline. Lipid desaturation enzymes such as FAT-7, the *C. elegans* homolog of mammalian SCD1, are transcriptionally upregulated in response to mating in the intestine and are required for maintaining maternal lipid reserves^112^. Because *ntr-1* and *ntr-2* are both downregulated by mating^112^, nematocin signaling is positioned as a candidate suppressor of this lipid remodeling response **(Figure 2B).**

**Figure 2:**
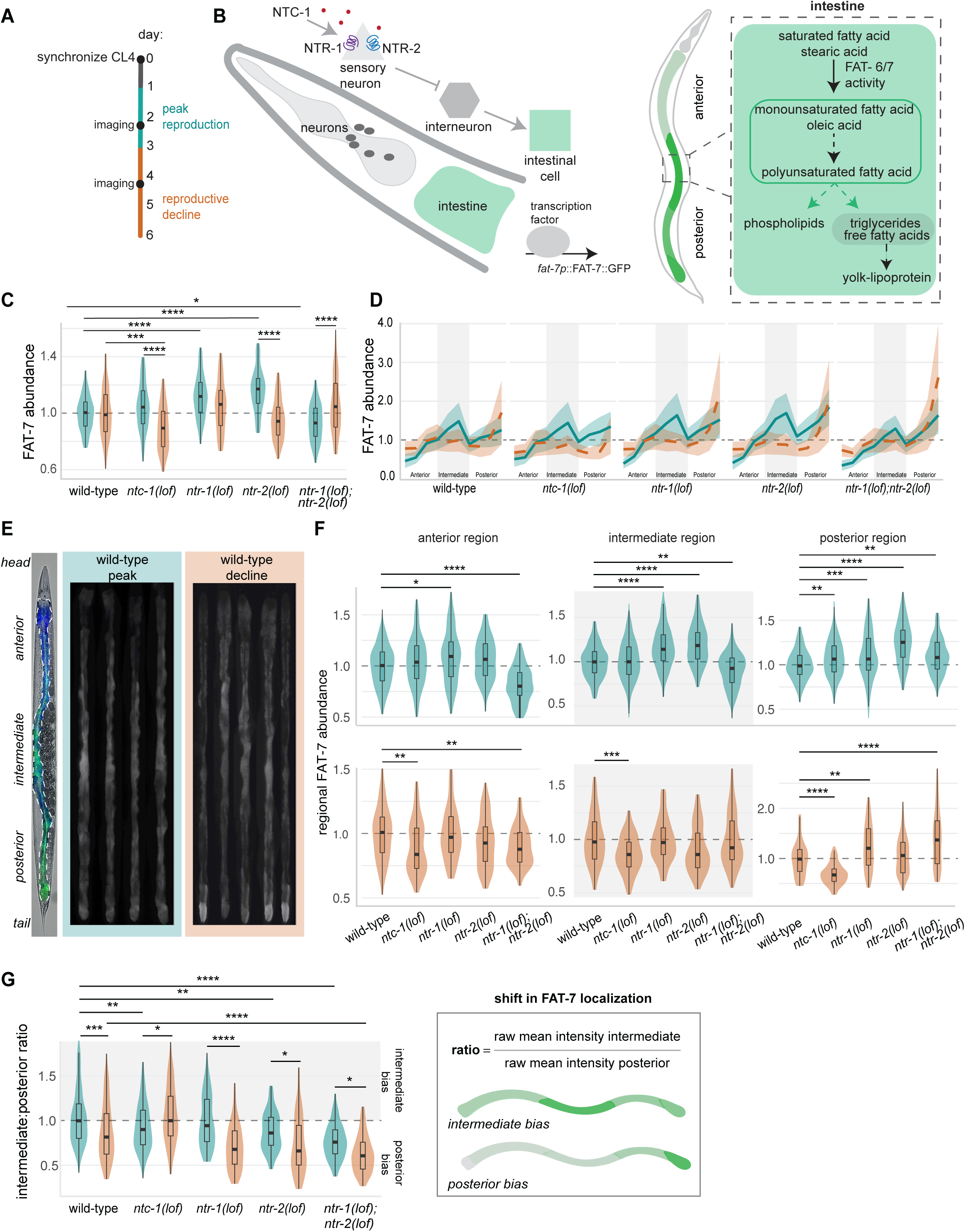
Loss of nematocin signaling is associated with altered intestinal FAT-7 abundance and regional distribution across reproductive age. **(A)** Experimental time course: All animals were synchronized with a 1–2-hour time egg lay, selected at crescent L4s and group housed at ∼ 50 worms per plate. Groups aged and transferred every other day to avoid starvation and crowding. Live imaging took place during peak reproduction (cyan) and the reproductive decline (orange), shown as black circles on the timeline. **(B)** Model of working hypothesis: Nematocin signaling suppresses intestinal FAT-7 abundance through though sensory neuron activity. Simplified metabolic pathway for fatty acid desaturation and yolk^2–4^. **(C)** Relative abundance of intestinal FAT-7 detected by fluorescence microscopy in *Pfat-7:fat-7::GFP* animals reared on OP50 until peak reproduction (adult day 2), or reproductive decline (adult day 4). All animals were normalized to their total intestinal area and corresponding wild-type replicate. **(D)** Relative abundance of FAT-7 across the segments of the intestine represented as a line plot where the mean (solid line) and standard error of the mean (ribbon) are shown per condition. All animals were normalized to their corresponding wild-type control. The segments are organized serially: anterior (1 - 3), intermediate (4 - 7) and posterior (8 - 10). **(E)** Representative images of wild-type intestines straightened by the WormToolbox module in CellProfiler at reproductive peak and decline. Dashed line on brightfield overlay represents the intestinal ROI straightened in the adjacent images. **(F)** Relative abundance of FAT-7 per region at each age. Intestinal segments were aggregated per animal, and each region was normalized to wild-type. The segments were combined into regions: anterior (1 - 3), intermediate (4 - 7) and posterior (8 - 10). **(G)** Intermediate:Posterior Ratio. The raw mean intensity of the intermediate region was divided by the mean intensity of the raw posterior region to evaluate the posterior shift in FAT-7 with reproductive age. Ratios greater than 1 represent animals with greater intermediate FAT-7 abundance relative to their posterior FAT-7 abundance. Values less than one are posterior biased. Diagram of the intestine illustrates intestinal FAT-7 localization and ratio calculation. **(C-F)** All conditions were normalized to wild-type mean. **(C-G)** 1 or 2-way ANOVA + TukeyHSD were applied. All relevant significant comparisons and adjusted p-values are shown. Adjusted *p*-values indicated by * *p* < 0.05, ** *p* <0.01, *** *p* < 0.001, **** *p* < 0.0001. Number of experimental and biological replicates: **(C)** Adult Day 2: N = 9, 9, 4, 3, 4; n = 263, 259, 166, 104,156; Adult Day 4: N = 4, 4, 4, 3, 4; n = 91, 85, 86, 72, 80 **(F)** Adult Day 2: N = 9, 9, 4, 3, 4, n = 200-203, 203-205, 109-112, 74-75, 101-103; Adult Day 4: N = 4, 4, 4, 3, 4; n = 87-92, 81-82, 83-85, 74-77, 77-80 **(G)** Adult Day 2: N = 9, 9, 4, 3, 4, n = 198, 196, 112, 72, 100; Adult Day 4: N = 4, 4, 4, 3, 4; n = 92, 82, 81, 75, 78

To test whether nematocin signaling suppresses intestinal lipid desaturation, we introduced our loss-of-function alleles into a FAT-7::GFP reporter^172^ and quantified total intestinal fluorescence at peak reproduction (adult day 2) and reproductive decline (adult day 4) (**Figure 2A, B**). Fluorescence intensity was normalized to intestinal area and to wild-type controls (**Figure 2C**).

During peak reproduction, loss of the peptide itself, *ntc-1,* was indistinguishable from wild type (1.00 vs 1.05; **Figure 2C**). By contrast, loss of either receptor alone increased intestinal FAT-7 abundance (1.00 vs 1.12, p < 0.0001 and 1.00 vs 1.16, p < 0.0001; **Figure 2C**), consistent with our prediction that nematocin signaling suppresses lipid desaturation. Unexpectedly, simultaneous loss of both receptors significantly reduced FAT-7 abundance relative to wild type (1.00 vs 0.94, p = 0 .03; **Figure 2C**) and loss of the peptide did not genocopy the double receptor mutant (1.05 vs 0.94, p < 0.0001; **Figure 2C**). This non-additive response suggests that NTR-1 and NTR-2 do not function redundantly but may engage distinct or antagonistic signaling states depending on receptor context.

During reproductive decline, the pattern shifted. Peptide loss now reduced FAT-7 abundance relative to wild type (1.00 vs 0.89, p = 0.0005; **Figure 2C**), and receptor loss no longer impacted total intestinal abundance (**Figure 2C**). Loss of *ntr-1* alone, however, did not display an age-dependent reversal in relative FAT-7 abundance (1.12 vs 1.05; **Figure 2C**).

In light of these findings, together with the age-dependent increase in *ntr-1* expression (**Figure 1A**), these data are consistent with nematocin signaling limiting intestinal FAT-7 abundance primarily through the dominant receptor NTR-1 during reproductive aging, with NTR-1 expression itself rising in an age-dependent manner that may amplify this effect over time.

#### FAT-7 localization is regionally regulated by nematocin receptor status

In previous studies, FAT-7 abundance has been quantified as total intestinal fluorescence^119,170,172–177^. However, the intestine is a structurally and functionally heterogeneous tissue. The anterior and intermediate intestine are the primary sites of nutrient absorption and active lipid metabolism, whereas the posterior intestine, adjacent to the rectum, is associated with defecation-cycle physiology and contains prominent neutral lipid storage pools^161,178–181^. Regional localization therefore distinguishes active lipid processing from lipid sequestration, especially in states of senescent obesity or steatosis.

To determine whether altered FAT-7 abundance **reflected enrichment** or **spatial redistribution**, we used the WormToolbox CellProfiler pipeline^182^ to segment each intestine into ten sequential anterior-to-posterior regions(**Figure 2D-E**) and quantified regional FAT-7 abundance per animal (**Figure 2F**) and localization bias (**Figure 2G**).

In wild-type animals, FAT-7 was broadly distributed during peak reproduction, with enrichment in the intermediate and posterior segments of the intestine (**Figure 2D & Supplementary Figure 5C**). By reproductive decline (day 4), this intermediate enrichment was substantially diminished, and FAT-7 became concentrated in the posterior intestine (**Supplementary Figure 5C**), a posterior bias not observed in young animals (1.04 vs 0.87, p < 0.0002; **Figure 2G**).

In *ntc-1*, *ntr-2* and *ntr-1;ntr-2* receptor mutants, the intermediate-to-posterior FAT-7 ratio was already shifted toward the posterior during peak reproduction before wild-type animals exhibited any such bias (1.04 vs 0.93, p = 0.006, 1.04 vs 0.88, p = 0.002, 1.04 vs 0.77, p <0.0001 respectively; **Figure 2G**).

This posterior enrichment became even more pronounced during reproductive decline in the double receptor mutants (0.87 vs 0.63, p < 0.0001 and 0.77 vs 0.63, p = 0.03; **Figure 2G**). Loss of *ntr-1* alone did not cause a posterior shift during peak reproduction (1.04 vs 0.99) and displayed a wild-type regional distribution with age (0.99 vs 0.70, p <0.0001; **Figure 2G**). By contrast, loss of the peptide, *ntc-1*, prevented the age-related redistribution. However, regional abundance was significantly lower at this time point across the board in *ntc-1(lof)* animals **(Figure 2F)**, suggesting that the absence of localization bias might be related to loss of signal overall. Collectively, the relative abundance and regional bias implies that peptide and receptor loss produce qualitatively distinct metabolic states.

The premature posterior bias observed in animals without *ntc-1* or *ntr-2* suggests that nematocin signaling normally restrains the posterior sequestration of FAT-7 during the reproductive period, possibly preserving intermediate intestinal lipid processing for fertility.

#### Loss of nematocin signaling enhances the mating-induced FAT-7 response

Because nematocin receptors are downregulated by mating and receptor mutants show elevated FAT-7 abundance, we next tested directly whether intact nematocin signaling restricts the metabolic response to mating.

We quantified intestinal FAT-7 abundance in unmated and mated wild-type and *ntc-1(lof)* animals during peak reproduction and normalized all measurements to unmated wild-type controls (**Figure 3A-C and Supplementary Figure 6**).

**Figure 3:**
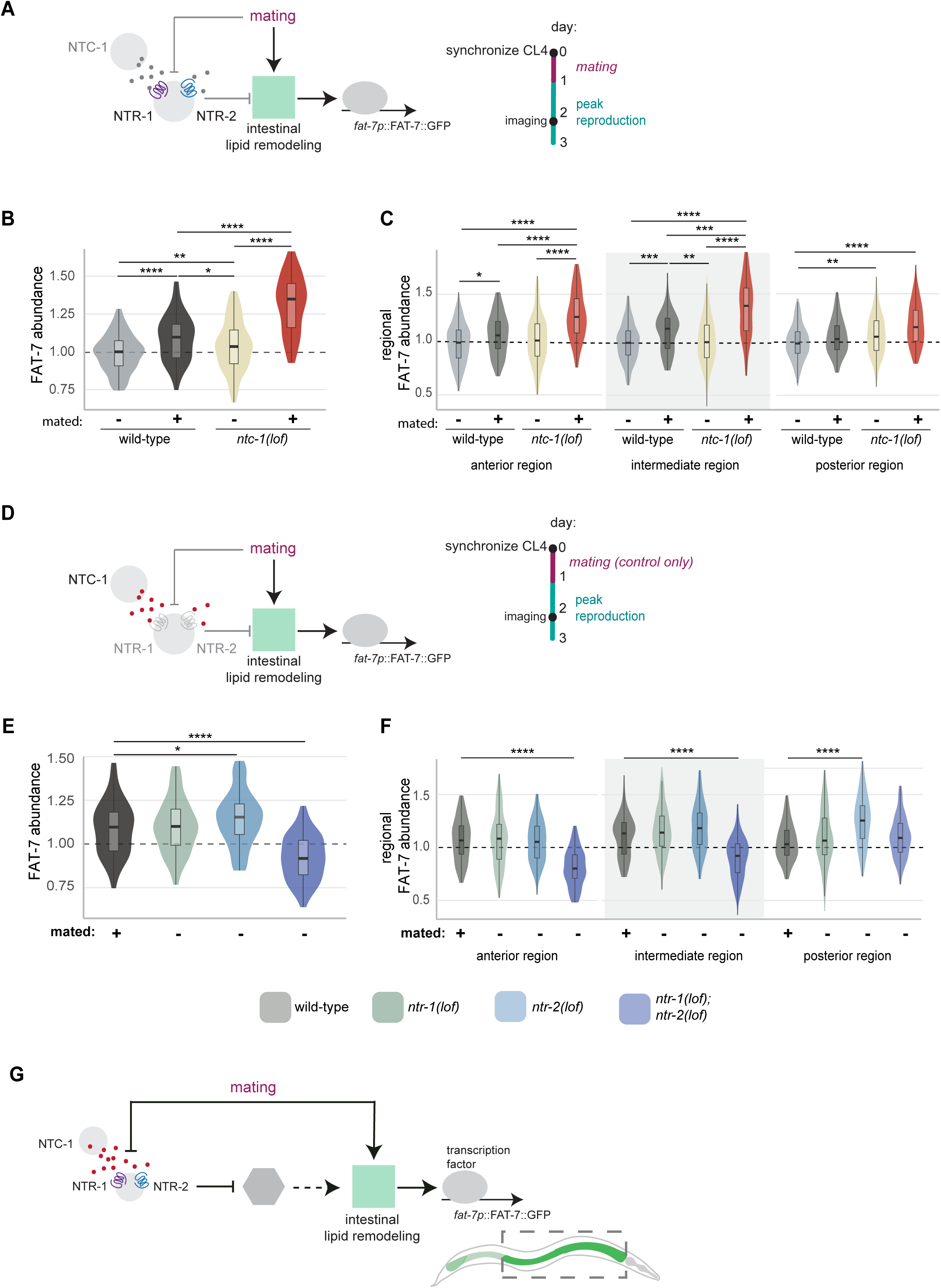
Nematocin signaling suppresses mating induced FAT-7 abundance. **(A)** Diagram of the working hypothesis. Nematocin and mating have opposing actions on intestinal FAT-7 abundance. Experimental timeline to the right. Unmated and mated animals were synchronized at crescent L4. Mated animals were paired with 3 wild-type young males for 24 hours and then group housed 20-50 animals per plate. Imaging took place ∼ 48 hours after synchronization. **(B)** Relative abundance of intestinal FAT-7 detected by fluorescence microscopy in unmated or mated *Pfat-7:fat-7::GFP* animals reared on OP50 until peak reproduction (adult day 2). All animals were normalized to their total intestinal area and corresponding unmated wild-type replicate. **(C)** Relative abundance of FAT-7 per region per condition. Intestinal segments were aggregated per animal and each region was normalized to unmated wild-type. **(D)** Diagram of working hypothesis. Nematocin receptor activity suppresses mating-induced FAT-7 abundance. Mated wild-type animals were compared to unmated receptor (lof) mutants. Experimental timeline to the right. Unmated and mated animals were synchronized at crescent L4. Mated wild-type animals were paired with 3 wild-type young males for 24 hours and then group housed 20-50 animals per plate. Imaging took place ∼48 hours after synchronization. **(E)** Relative abundance of intestinal FAT-7 detected by fluorescence microscopy in unmated or mated *Pfat-7:fat-7::GFP* animals reared on OP50 until peak reproduction, adult day 2. All animals were normalized to their total intestinal area and corresponding unmated wild-type replicate. **(F)** Relative abundance of FAT-7 per region per condition. Intestinal segments were aggregated per animal and each region was normalized to unmated wild-type. **(B/E)** All animals were normalized to the wild-type median. Dunn Test + Benjamini-Hochberg (FDR) corrections were applied to pooled samples between all genotypes **(C/F)** All animals were normalized to the wild-type mean. 1-way ANOVA + TukeyHSD were applied. All relevant significant comparisons and adjusted p-values are shown. Adjusted *p*-values indicated by * *p* < 0.05, ** *p* <0.01, *** *p* < 0.001, **** *p* < 0.0001. Number of experimental and biological replicates: **(B)** N = 9, 4, 9, 4; n = 263, 83, 259, 80 **(C)** N = 9, 4, 9, 4; n = 200-203, 71, 202-205, 67-68 **(E)** N = 3, 4, 3, 4; n = 83, 166, 104, 156 **(F)** N =9, 4, 3, 4; n = 71, 109-112, 74-75, 101-103

Mating increased total intestinal FAT-7 by approximately 10% in wild-type animals (1.00 vs 1.10, p < 0.0001; **Figure 3B**). In *ntc-1(lof)* animals, the same mating event elevated FAT-7 by approximately 35% (1.00 vs 1.35, p < 0.0001; **Figure 3B**), indicating that intact nematocin signaling suppresses nearly 70% of the normal mating-induced FAT-7 response (1.10 vs 1.35, p < 0.0001; **Figure 3B**). Importantly, mated *ntc-1(lof)* animals reached FAT-7 levels substantially above those observed in unmated receptor mutants (1.35 vs 1.10, p < 0.0001; 1.35 vs 1.15, p < 0.0005; 1.35 vs 0.92, p < 0.0001; **Supplementary Figure 7**), indicating that mating and receptor loss do not simply saturate the same pathway.

Regional analysis identified the intermediate intestine as the primary site of this response (**Supplementary Figure 6**). Mating increased FAT-7 abundance in the anterior and intermediate intestine (1.00 vs 1.10, p = 0.005 and 1.00 vs 1.10, p = 0.0008; **Figure 3C**), but the increase was substantially larger in *ntc-1(lof)* animals (1.00 vs 1.29, p < 0.0001 and 1.00 vs 1.33, p < 0.0001; **Figure 3C**). By contrast, the posterior intestine was not significantly affected by mating in wild-type animals (1.00 vs 1.06) and modestly increased in *ntc-1(lof)* (1.00 vs 1.19, p < 0.0001 and 1.07 vs 1.19, p = 0.002; **Figure 3C**).

Interestingly, the intermediate intestine is also the region where FAT-7 abundance declines most strongly during age-dependent redistribution in unmated wild-type animals (1.03 vs 0.87, p = 0.0002; **Figure 2G**). This suggests that reproductive status and aging regulate the same metabolic compartment. Taken together, these data provide direct evidence that intact nematocin signaling suppresses the mated FAT-7 response by limiting its enrichment to the intermediate intestine.

#### Unmated receptor mutants partially phenocopy the mated wild-type FAT-7 state through region-specific intestinal remodeling

Although mating was required to fully saturate the FAT-7 response in *ntc-1(lof)* animals, we next asked whether unmated receptor mutants already resembled the mated wild-type state (**Figure 3D-F & Supplementary Figure 7A-B**). If nematocin normally suppresses mating-induced lipid remodeling, loss of receptor function in unmated animals could create a constitutively “primed” metabolic state that is partially phenocopied by mating.

Total intestinal FAT-7 abundance was not significantly different between mated wild-type animals and either single receptor mutant, *ntr-1(lof)* or *ntr-2(lof)*, in the unmated state (1.10 vs 1.10, and 1.10 vs 1.15; **Figure 3E**), consistent with a primed-response model. In contrast, animals lacking both receptors had significantly lower FAT-7 levels than mated wild-type controls (1.10 vs 0.92, p <0.0001; **Figure 3E**), indicating that the double mutant does not simply represent a constitutively mated metabolic state.

We next asked whether this putative priming was reflected in regional FAT-7 distribution, particularly in the intermediate intestine where mating produced the strongest increase (**Figure 3**). The intermediate region was significantly enriched in both *ntr-1(lof)* and *ntr-2(lof)* animals relative to unmated wild-type controls (**Figure 3**) but was not different from mated wild-type animals (1.10 vs 1.16 and 1.10 vs 1.19; **Figure 3F**), further supporting a primed physiological state. By contrast, the posterior intestine was unaffected by mating in wild-type animals (1.00 vs 1.06; **Figure 3C**), yet all receptor mutant combinations showed elevated posterior FAT-7 abundance relative to unmated wild type (**Figure 2E**) but were not different from mated *ntc-1(lof)* (**Supplementary Figure 7B**).

Together, these data support two separable modes of nematocin regulation. One promotes FAT-7 enrichment in the intermediate intestine in response to reproductive status (NTR-1 or NTR-2) and mating (NTC-1), while the other restrains posterior intestinal FAT-7 accumulation independently of age (NTR-2).

### Nematocin signaling is associated with altered yolk lipoprotein abundance in the intestine and oocyte in a receptor- and reproductive-age-dependent manner

#### Intestinal VIT-2 abundance in nematocin receptor mutants shows an age-dependent reversal

FAT-7 abundance prepares the intestine for lipid desaturation, but the reproductive consequence of this metabolic state depends on how those lipids are ultimately used. A large fraction of desaturated fatty acids are incorporated into triglycerides that support yolk synthesis, making intestinal yolk production a major downstream consequence of lipid remodeling^142,179,183^.

Because oocyte quality^152^ and fertilization efficiency depend on maternal provisioning^138,139,183^, we next asked whether altered FAT-7 abundance was reflected in changes in yolk lipoprotein abundance^112,175^ . To detect changes in metabolic investment toward reproduction, we measured VIT-2::GFP abundance at the site of synthesis (intestine) and delivery (oocyte) during peak reproduction (day 2) and reproductive decline (day 4) (**Figure 4A-B**).

**Figure 4:**
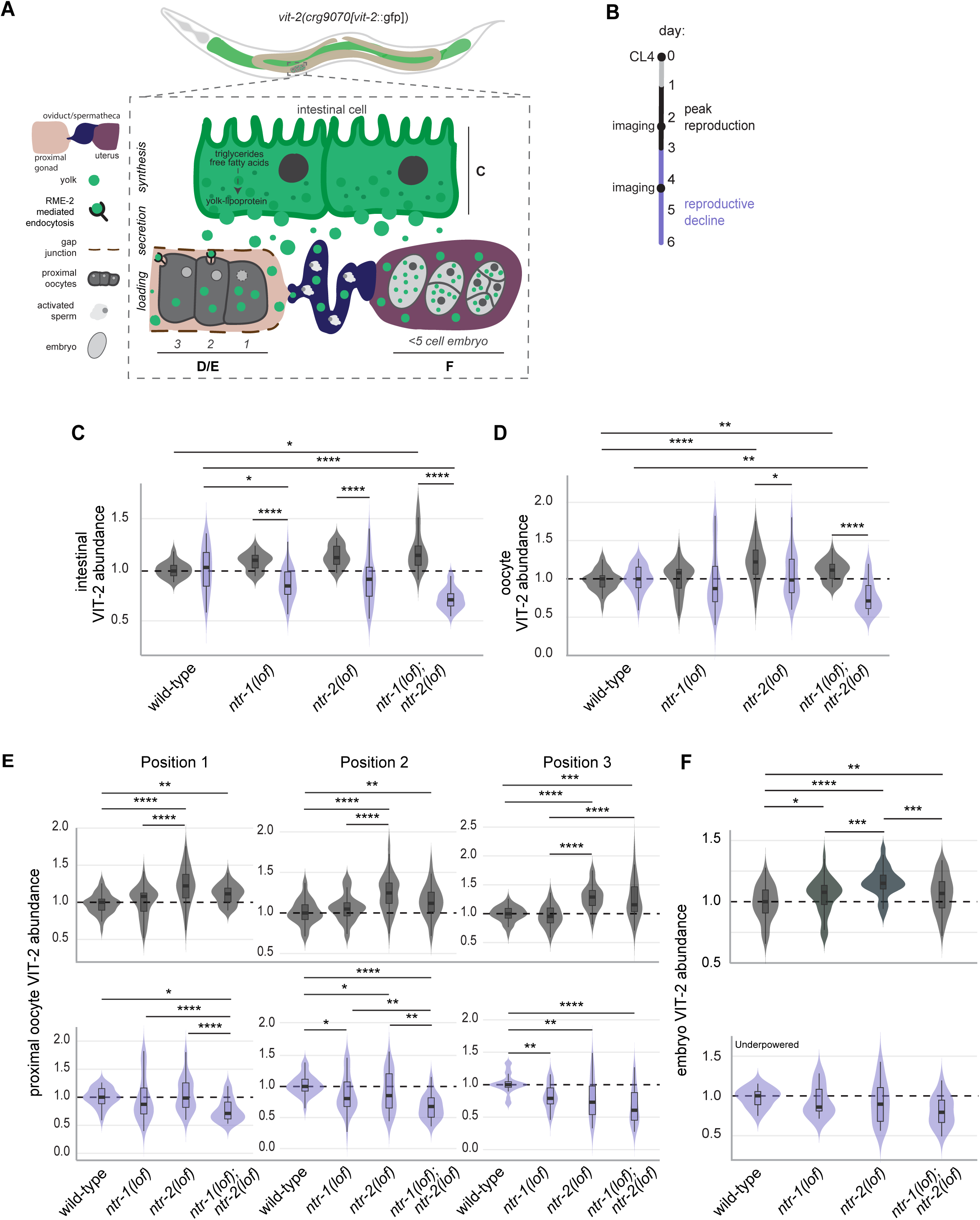
Nematocin signaling preserves nutritional provisioning to the egg with age at the expense of peak reproductive investment. **(A)** Diagram of intestinal yolk secretion and loading into the germline through receptor-mediated endocytosis by RME-2 with designated figure annotations. **(B)** Experimental time course: All animals were synchronized with a 1–2-hour time egg lay, selected at crescent L4s and group housed at ∼ 50 worms per plate. Groups aged and transferred every other day to avoid starvation and crowding. Live imaging took place during peak reproduction (black) and the reproductive decline (purple). **(C)** Relative abundance of intestinal VIT-2 detected by fluorescence microscopy in *Pvit-2:vit-2::GFP* animals reared on OP50 until peak reproduction (adult day 2) or reproductive decline (adult day 4). All animals were normalized to their total intestinal area and corresponding wild-type replicate. **(D)** Relative abundance of VIT-2 in position 1 oocytes detected by fluorescence microscopy in *Pvit-2:vit-2::GFP* animals reared on OP50 until peak reproduction (adult day 2) or reproductive decline (adult day 4). All intensity measurements were divided by their oocyte area, averaged per animal and normalized to wild-type. **(E)** Relative median abundance of VIT-2 in proximal oocytes at positions 1-3 relative to the spermatheca/oviduct. **(F)** Relative abundance of VIT-2 in early embryos (< 5-cell) and eggs (1-cell) during reproductive peak or decline. **(B/D)** All samples were normalized to wild-type mean. 2-way ANOVA + TukeyHSD were applied. All significant comparisons and adjusted p-values are shown. Adjusted *p*-values indicated by * *p* < 0.05, ** *p* <0.01, *** *p* < 0.001, **** *p* < 0.0001. Number of experimental and biological replicates: **(B)** Adult Day 2: N = 2, 2, 2, 2; n = 18, 23, 30, 20 and Adult Day 4: N =3, 3, 3, 3; n = 28, 35, 36, 23) **(C)** Adult Day 2: N = 3, 3, 3, 3; n = 42, 42, 47, 45 and Adult Day 4: N = 3, 3, 3, 3; n= 32, 35, 34, 29 **(E/F)** All samples were normalized the wild-type median. Dunn Test + Benjamini-Hochberg (FDR) corrections were applied to pooled samples between all genotypes. All significant comparisons and adjusted p-values are shown. Adjusted *p*-values indicated by * *p* < 0.05, ** *p* <0.01, *** *p* < 0.001, **** *p* < 0.0001. Number of experimental and biological replicates: **(E)** Adult Day 2: N = 3, 3, 3, 3; n = 36-42, 33-43, 37-47, 32-34 and Adult Day 4: N = 3, 3, 3, 3; n = 28-34, 31-35, 35-36, 29-30 **(F)** Adult Day 2: N = 4, 4, 4, 4; n = 63, 62, 61, 67 and Adult Day 4: N = 1, 1, 1, 1; n = 12, 16, 18, 18

To determine whether nematocin signaling supports yolk production, we crossed receptor mutants into an endogenous VIT-2::GFP reporter^184^ and quantified mean fluorescence in the intestine, normalized to the median wild-type control (**Figure 4C**).

If FAT-7 abundance directly predicted provisioning, then, during peak reproduction, we expected increased intestinal yolk in single receptor mutants and reduced abundance in the double mutant. Instead, we observed the opposite pattern: intestinal VIT-2 was significantly increased in the double receptor mutant (1 vs 1.18; p = 0.02; **Figure 4C**), while single receptor mutants showed little or no change (+ 8-13%).

During reproductive decline however, this relationship reversed. Loss of *ntr-1* alone or in combination with *ntr-2,* significantly reduced intestinal VIT-2 relative to wild type (1.00 vs 0.86, p = 0.02, and 1.00 vs 0.72, p < 0.0001; **Figure 4C**). Comparing young and old animals revealed that loss of either receptor or both in combination reversed the mutant relationship to wild type across age (1.08 vs 0.87, p < 0.0001, 1.13 vs 0.89, p < 0.0001 and 1.18 vs 0.72, p <0.0001; **Figure 4C**).

These findings suggest that nematocin signaling mediates an age-dependent tradeoff: modestly restraining early-life yolk abundance while supporting maintenance of intestinal yolk stores during reproductive decline. This temporal shift is not fully explained by FAT-7 abundance alone, indicating that yolk production reflects both lipogenesis and additional regulation of secretion and transport.

#### Loss of *ntr-2* improves yolk loading efficiency into the egg

Because intestinal VIT-2 abundance depends on both synthesis and secretion, increased intestinal yolk does not necessarily indicate improved provisioning to the embryo. Efficient maternal investment also requires uptake into the oocyte through RME-2-mediated receptor endocytosis^147^.

To determine whether nematocin signaling affected nutritional provisioning to the egg, we measured VIT-2::GFP abundance in the three oocytes most proximal to the spermatheca/oviduct (positions 1–3), where final yolk loading occurs prior to ovulation (**Figure 4A, D-F**).

During peak reproduction, loss of *ntr-2* alone or in combination with *ntr-1* increased yolk abundance in position 1 oocytes, which represent the final stage before ovulation, compared to wild type (0.97 vs 1.22, p < 0.0001; **Figure 4D**), despite only a modest increase in intestinal VIT-2 in *ntr-2(lof)* animals. This disproportionate increase suggested loss of *ntr-2* also improves yolk loading efficiency rather than increasing production alone.

To test this directly, we asked whether yolk loading was increased across all positions. *ntr-2(lof)* animals showed approximately 25% greater yolk abundance at each successive oocyte position (1.00 vs 1.25, p < 0.0001 and 1.00 vs 1.29, p < 0.0001; **Figure 4A, E**), consistent with sustained enhancement of loading throughout late oogenesis. We further validated this phenotype in early embryos containing fewer than five cell divisions, confirming elevated yolk content at the time of fertilization (1.00 vs 1.15, p < 0.0001; **Figure 4A, F**).

Loss of both receptors increased oocyte yolk by 11% (1.00 vs 1.11, p = 0.003; **Figure 4E**), which was similar to loss of either receptor (1.08 vs 1.11; p = 0.07 and 1.11 vs 1.22, p = 0.07; **Figure 4E**). While the double mutant was not statistically different from *ntr-2(lof)*, the loss of *ntr-1* reduced the yolk content by half. This suggests that signaling through NTR-1 may be required to increase yolk loading efficiency in the germline of *ntr-2(lof)* animals. This finding was corroborated in early embryos where elevated yolk was detected in all mutant combinations and the differences between the single and double receptor mutants were clear **(Figure 4F)**.

Together, these data support the hypothesis that loss of *ntr-2* improves yolk loading efficiency in the oocyte, likely through enhanced RME-2-dependent uptake. These findings suggest receptor specialization within the nematocin pathway, where NTR-1 primarily regulates intestinal metabolic state and reproductive output, while NTR-2 more strongly influences yolk uptake by the germline.

#### During reproductive decline, double receptor mutants become substrate-limited

During reproductive decline, the phenotype shifted substantially. *ntr-1;ntr-2(lof)* animals showed reduced VIT-2 abundance in both the intestine and proximal oocytes, including a marked reduction in position 1 oocytes (1.00 vs 0.76, p = 0.003; **Figures 4C-E**) and across all subsequent loading stages (1.00 vs 0.68, p < 0.0001 and 1 vs 0.61, p < 0.0001; **Figure 4E**). This phenotype paralleled the reduction in intestinal yolk stores, suggesting that reduced substrate availability, not impaired uptake, was the primary limiting factor.

Loss of either receptor alone, *ntr-1* or *ntr-2*, did not significantly reduce yolk abundance in the final pre-ovulatory oocyte (1.00 vs 0.97 and 1.00 vs 1.04; **Figure 4D**), despite reductions at position 2 (1.00 vs 0.81 (*ntr-1*), p = 0.02 and 1.00 vs 0.79 (*ntr-2*), p = 0.005; **Figure 4E**) and position 3 (1.00 vs 0.85 (*ntr-1*), p = 0.04 and 1.00 vs 0.73 (*ntr-2*), p = 0.002; **Figure 4E**). This indicates that enhanced yolk loading remains sufficient to restore normal provisioning before ovulation when intestinal supply is only modestly reduced.

Thus, yolk loading is preserved unless intestinal substrate availability becomes limiting. Only the double mutant crosses this threshold during reproductive decline.

Taken together, these findings support a model in which nematocin signaling restrains maternal metabolic investment during reproductive aging by suppressing intestinal lipid desaturation and yolk loading into the oocyte. Loss of this suppression improves mid-life fertility by enhancing both oocyte quality and reproductive output. NTR-1 primarily regulates intestinal FAT-7 abundance and reproductive output, whereas NTR-2 acts at the oocyte to influence yolk loading efficiency. These results establish nematocin as a neuroendocrine regulator of maternal investment across the reproductive lifespan.

## Discussion

### Nematocin signaling limits reproductive capacity during reproductive aging

Reproductive aging is often described as the progressive depletion and deterioration of gametes. While germline decline is undeniably central to reproductive senescence, increasing evidence across taxa suggests that reproductive aging is also shaped by systemic physiology, particularly by neuroendocrine pathways that regulate the allocation of somatic resources to reproduction. Here, we identify the oxytocin/vasopressin-like neuropeptide system nematocin as a regulator of maternal metabolic investment during reproductive aging in *C. elegans*.

We found that expression of *ntc-1*, *ntr-1*, and *ntr-2* remained elevated throughout reproductive adulthood, peaking during the onset of reproductive decline rather than during the period of maximal oocyte production (**Figure 1**). Consistent with this expression pattern, loss of nematocin signaling improved mated fecundity in reproductively middle-aged animals, with the strongest effects observed in *ntr-1* and *ntr-1;ntr-2* mutants. Increased reproductive capacity reflected both enhanced ovulation and improved fertilization efficiency, whereas embryonic viability remained unchanged. Because embryo survival was unaffected, the fertility advantage likely arises prior to embryogenesis and is most consistent with improvements in oocyte quality rather than post-fertilization rescue.

Together, these findings identify nematocin signaling as a regulator of reproductive capacity during mid-life. Unlike many interventions, loss of nematocin signaling improves fertility during reproductive decline without an obvious reduction in early reproductive performance (**Supplementary Figure 1**). This interpretation is further supported by the temporal pattern of pathway expression. If nematocin primarily functioned to promote reproduction, maximal signaling would be expected during peak reproductive output. Instead, expression peaks during the onset of reproductive decline, consistent with a role in limiting reproductive investment as animals age. We therefore propose that nematocin signaling acts as a biological brake on reproductive capacity during mid-life.

### Improved fertilization efficiency suggests enhanced oocyte quality

Fertilization efficiency reflects many aspects of oocyte quality, including those that support sperm migration and guidance^152,153,153–156,185^. In *C. elegans*, oocyte lipid composition is largely determined by intestinal nutrient provisioning. Polyunsaturated fatty acids (PUFAs), delivered to developing oocytes through yolk lipoproteins^138^, serve as substrates for prostaglandin synthesis that promotes sperm motility and chemotaxis^138,139,183,186,187^. Disruption of PUFA availability, whether through dietary manipulation or altered neuroendocrine signaling^139,163,185^, impairs sperm guidance and reduces fertilization efficiency.

These findings suggest that at least part of reproductive decline may reflect age-associated changes in maternal provisioning^157,184,188–190^ rather than irreversible deterioration of the oocyte itself. In this view, oocyte quality remains responsive to extra-gonadal physiology and environmental signals.

Although we observed increased fertilization efficiency in *ntr-1* and *ntr-1;ntr-2* mutants, we did not directly measure sperm migration, prostaglandin production, or oocyte fatty acid composition.

Consequently, further work is needed to identify the mechanisms by which altered nutritional provisioning leads to the fertility phenotypes observed here. Nevertheless, the improvement in fertilization efficiency is consistent with a model in which loss of nematocin signaling enhances oocyte quality through changes in maternal lipid metabolism.

### Nematocin does not appear to enhance fertility through maintenance of the germline progenitor pool

In addition to increased fertilizability, loss of nematocin signaling increased overall reproductive output. Reproductive capacity in aging *C. elegans* is strongly influenced by the maintenance of germline stem cell activity and the rate at which progenitor cells enter meiosis^191,192^. Notably, relatively few interventions increase reproductive output during reproductive decline without substantially compromising early-life reproduction. For example, disruption of *che-3*^193^ or overexpression of *sygl-1* prolongs reproductive capacity by maintaining the progenitor zone and increasing meiotic output in mid-life^192^. In contrast, longevity-associated interventions such as reduced insulin signaling or dietary restriction often preserve late-life reproductive potential at the expense of early reproductive performance^6,132,133^ by maintaining the progenitor zone but also suppressing meiotic entry^192,194,195^.

Sensory perception also influences germline maintenance during reproductive aging. Exposure to male pheromones expands the progenitor zone and delays reproductive decline through signaling pathways involving serotonergic neurons (NSM, HSN), ASI sensory neurons, and the germline via DAF-7/TGFβ signaling^123,196^. An additional unknown signal from the ADL sensory neurons is required for this response^197^. Notably, both *ntr-1* and *ntr-2* are expressed in ADL neurons and are co-expressed in several *che-3*-positive sensory neurons (**Supplementary Figure 4**), supporting the possibility that nematocin signaling influences reproductive output through sensory regulation of germline activity.

However, we did not detect an expansion of the progenitor zone in nematocin signaling mutants (**Supplementary Figure 3**). This suggests that the increase in reproductive output either occurs downstream of stem cell maintenance or reflects changes too subtle to detect through progenitor-zone row measurements alone. Because meiotic entry was not assessed directly, we cannot exclude the possibility that nematocin signaling influences the transition from progenitor cell to developing gamete. Future studies examining meiotic progression and germline dynamics will be necessary to distinguish between these possibilities.

### Nematocin couples reproductive status to intestinal metabolism

*ntc-1* is expressed by AFD and AWC neurons, while the receptors *ntr-1* and *ntr-2* are expressed in ADF, ADL, and ASK sensory neurons that communicate with peripheral metabolic tissues and integrate social and mating cues. Thus, our data position nematocin within a broader neuroendocrine network coupling reproductive status to systemic metabolism.

In *C. elegans*, the intestine is particularly well suited for this role because it performs the combined functions of mammalian liver and adipose tissue while also serving as the site of vitellogenin synthesis. Maternal provisioning in this system is therefore fundamentally an intestinal process, linking somatic metabolic state directly to fertility.

### Nematocin suppresses mating-induced intestinal lipid remodeling

Our findings support a model in which nematocin signaling limits nutritional provisioning to the germline by suppressing yolk production early in reproduction and intestinal lipid desaturation with age, thereby preserving intestinal metabolic capacity for continued provisioning to developing oocytes. This model is further supported by the observation that intact nematocin signaling restrains mating-induced lipid remodeling and mid-life mated fertility.

Our analysis of FAT-7 supports the intestine as a major site of nematocin effects (**Figure 2**). FAT-7, the stearoyl-CoA desaturase homologous to mammalian SCD1, is a central regulator of monounsaturated fatty acid production and a key component of mating-induced lipid remodeling^112^. Loss of *ntr-1* consistently increased intestinal FAT-7 abundance across reproductive age, while mating induced a substantially larger FAT-7 response in *ntc-1* mutants than in wild type. These findings provide direct evidence that intact nematocin signaling suppresses the metabolic response to mating (**Figure 3**). The reduction of FAT-7 below wild-type levels in *ntr-1;ntr-2(lof)* during peak reproduction (**Figure 2C**) suggests that NTR-1 and NTR-1 are not simply redundant.

Because mating normally downregulates *ntr-1* and *ntr-2*^112^, receptor loss may partially mimic a constitutively post-mating metabolic state that primes animals for enhanced reproductive output. Consistent with this, unmated single receptor mutants showed intermediate intestinal FAT-7 abundance comparable to mated wild-type levels (**Figure 3E, F**).

### Reproductive aging is accompanied by regional reorganization of intestinal lipid metabolism

A second, less expected feature of our data is the spatial reorganization of intestinal FAT-7 that accompanies reproductive aging (**Figure 2**). In wild-type animals, FAT-7 shifts from modest enrichment in the intermediate intestine during peak reproduction to pronounced posterior accumulation during reproductive decline.

Regional analysis further refined this model. The intermediate intestine is the primary site of active lipid absorption and processing, whereas the posterior intestine is associated with lipid storage, temperature sensing, and defecation-cycle physiology^134,146,178^. Premature posterior bias in nematocin receptor mutants suggests that nematocin normally restrains posterior sequestration of FAT-7 and helps preserve metabolically productive intermediate intestinal states during the reproductive period.

The biological significance of this shift is not immediately obvious. It may reflect active sequestration of desaturase activity into compartments specialized for lipid storage; depletion of intermediate-region FAT-7 through sustained consumption for substrate production; or a more general age-associated change in the metabolic character of anterior and intermediate intestinal cells.

Several lines of evidence indicate that regional reorganization is a general feature of how the *C. elegans* intestine remodels its metabolic state. Posterior-biased lipid signal has been noted under social^114,171^, dietary^173,174,198^, thermal^170,172,177^, and fasting perturbations^199,200^. Most directly relevant, Goncalves et al. described a posterior shift in FAT-6 in aging males and linked this redistribution to reproductive tissues^201^. Our data extend this principle to hermaphrodites and suggest that reproductive cues actively maintain intermediate-intestinal desaturase capacity during periods of maximal reproductive demand.

Together, these observations identify specific intestinal regions as operative units of metabolic regulation during reproductive aging.

### Nematocin regulates maternal provisioning through distinct effects on lipid remodeling and yolk transfer

Because desaturated fatty acids are incorporated into triglycerides that support vitellogenin synthesis^142^, FAT-7 abundance provides an upstream indicator of provisioning capacity, whereas VIT-2 abundance reflects the downstream execution of maternal investment(**Figure 4**).

Intestinal VIT-2 abundance showed an age-dependent reversal across receptor mutants: during peak reproduction, the double receptor mutant accumulated excess intestinal yolk, whereas during reproductive decline, loss of *ntr-1* alone or in combination with *ntr-2* reduced intestinal yolk stores. Yolk production generally reflected yolk loading into the germline in all mutants except *ntr-2*, which transferred more yolk into oocytes relative to intestinal abundance.

These findings, together with the FAT-7 results, support a model in which NTR-1 primarily regulates the intestinal metabolic environment, whereas NTR-2 more strongly influences how efficiently maternal resources are transferred to developing oocytes. This suggests that nematocin signaling mediates a temporal tradeoff in which early restraint of provisioning supports maintenance of intestinal reserves later in life.

One interesting feature of our data is that intestinal FAT-7 and oocyte yolk abundance do not always track each other in time. Several factors plausibly contribute to this temporal mismatch. FAT-7 abundance reflects desaturase capacity rather than metabolic flux. Desaturated fatty acids also have multiple downstream fates, including incorporation into triglycerides for vitellogenin assembly, storage in lipid droplets, conversion into longer-chain PUFAs, or oxidation for energy production.

In addition, yolk loading depends not only on intestinal vitellogenin synthesis but also on secretion and oocyte uptake through RME-2-mediated endocytosis^147^. The temporal mismatch between FAT-7 and yolk therefore does not contradict the model but rather highlights the multiple regulatory steps linking intestinal lipid metabolism to effective maternal provisioning.

### Maternal provisioning depends on both lipid quantity and lipid quality

Insufficient yolk provisioning has well-established consequences for fertilization and offspring fitness, including impaired sperm guidance^138,139^, delayed larval development^141^, reduced starvation resistance^144,184^, and increased susceptibility to starvation-induced reproductive abnormalities^144^.

However, the relationship between yolk abundance and offspring quality is not strictly proportional. Additional yolk does not necessarily confer proportional improvements in embryonic or larval fitness under favorable conditions^184^. Evidence from previous studies suggests that benefits of yolk provisioning may plateau during peak reproduction, whereas additional provisioning may become advantageous under future nutrient limitation^144^.

Importantly, yolk production is metabolically costly. Elevated yolk synthesis contributes to age-associated somatic lipid depletion^137^ and intestinal atrophy^134,135^, highlighting a trade-off between maternal maintenance and offspring provisioning. Consistent with this idea, oleic acid supplementation protects against age-induced somatic fat loss^137^, mating-induced lipid depletion^112^, and mating-induced lifespan shortening^112^ while enhancing embryonic stress resistance^143,171^. However, oleic acid supplementation does not increase mated progeny production but does rescue fecundity under nutrient stress^175^.

Our findings show that loss of nematocin signaling increases yolk loading by approximately 8–22% during peak reproduction, a range comparable to that observed during maternal aging^184^ and dietary restriction^144^. Whether this increase benefits offspring fitness or imposes costs on maternal physiology remains untested.

Together, these observations suggest that reproductive success may depend not only on the quantity of lipids transferred to oocytes but also on their composition. Although yolk abundance is commonly used as a proxy for maternal provisioning, total lipid measurements do not capture differences in lipid quality^138^.

Recent work has demonstrated that pheromone exposure remodels intestinal lipid metabolism through NHR-49, increasing FAT-7 expression, promoting lipid mobilization, and enriching oocytes with PUFAs despite no measurable increase in yolk loading^171^. Thus, changes in oocyte quality may reflect alterations in lipid composition rather than total yolk content. This may explain why FAT-7 expression does not consistently correlate with yolk abundance in our experiments yet remains associated with NTR-1’s fertility outcomes. If elevated fertility is driven by increased PUFA availability downstream of FAT-7 activity, then oocytes may acquire a more favorable lipid composition without necessarily accumulating more yolk.

### A sensory-neuroendocrine model for reproductive investment during aging

Notably, pheromone-induced remodeling of oocyte lipid composition requires signaling through ASK sensory neurons^171^, which express NTR-1. GPCR signaling within ASK neurons is necessary for this response, suggesting a neuronal mechanism linking environmental cues to maternal lipid allocation.

If ASK neurons also contribute to the FAT-7 changes observed here, NTR-1 signaling could act to suppress ASK activity, thereby limiting intestinal lipid remodeling required for enhanced oocyte PUFA enrichment and its downstream benefits. Consistent with a role for ASK in regulating lipid metabolism, ASK neurons, together with ASG, AWB, AWC, ASI, and ASJ neurons, have been implicated in suppression of FAT-7 expression during neuronal temperature stress^170^.

### Parallels to mammalian lactation

Maternal transfer of energetic resources begins at oogenesis and extends through adulthood. Across species, the maternal metabolic state must not only balance current and future reproductive fitness but also support multiple stages of offspring development.

While vitellogenin genes carry out the bulk of maternal provisioning in *C. elegans*^136^, their vertebrate homologues, very low-density lipoproteins (VLDLs)^202^, mirror that function in mammals^203^. Secreted by the liver, VLDLs transport essential nutrients to the egg in egg-laying animals. In addition to hepatic VLDLs, mammalian lactation also utilizes lipids transferred from the small intestine (chylomicrons)^204,205^ and adipose tissue (free fatty acids bound to albumin)^206^.

The mammalian oxytocin receptor, OXTR, is expressed in many metabolic tissues^91,97,100,207–210^ and is known to centrally regulate feeding behavior^72^, promote energy expenditure^211^ and support various reproductive processes including milk ejection^68^ during lactation. Until recently, the tissue specific contributions of oxytocin’s actions on peripheral metabolism had not been elucidated.

Li et al., 2024 identified sympathetic release of oxytocin in adipose tissue as a necessary enhancer of lipolysis^209^ upon stress, fasting, and cold exposure. In 2026, they expanded their model to include lactation^210^. The mammary gland requires circulating free fatty acids liberated from adipose to synthesize triglyceride rich milk. Under standard feeding conditions (16.5% fat) OXTR signaling is required to enhance adipocyte lipolysis during lactation. Although, feeding adipocyte-specific OXTR knockout dams a high fat diet (60% fat) rescued milk composition and pup growth during lactation. This finding demonstrates that oxytocin signaling mobilizes maternal lipid reserves to offspring depending on maternal energy state.

Our findings support a model where nematocin promotes maternal lipid transfer with age. In the absence of nematocin signaling, yolk production and secretion is reduced over time, suggesting that nematocin signaling mutants may be sequestering lipids as animals lose metabolic flexibility with age. This finding parallels the OXTR adipocyte-mammary axis in mammals, where nematocin signaling regulates provisioning depending on maternal energy balance and is required to maximize lipid transfer during maternal lipid depletion.

Collectively, these findings support a model in which NTR-1-dependent sensory signaling restrains intestinal lipid desaturation and reproductive output. We propose that nematocin integrates sensory and metabolic information to modulate reproductive investment as animals age. Through this neuroendocrine pathway, environmental and social cues may influence intestinal lipid remodeling, oocyte provisioning, and fertility during reproductive decline.

Future studies combining tissue-specific receptor manipulations, lipidomic analysis, prostaglandin measurements, and direct examination of germline dynamics will be necessary to fully define how nematocin signaling coordinates metabolism and reproduction during aging.

## ACKNOWLEDGEMENTS

We thank members of the Garrison lab for constructive discussions and Frontiers in Reproductive: Molecular and Cellular Concepts and Applications training course, Marissa Dobry, and Bikem Soygur for technical assistance; the Caenorhabditis Genetics Center (CGC), which is funded by the NIH Office of Research Infrastructure Programs (P40 OD010440), the National Bioresource Project, and for strains. This work was funded in part by the NIH (R35 GM119828, R35 GM145305, and S10 OD021686 to J.L.G., NIA T32 AG052374 to K.M.A.) and the Juno Institute.

## AUTHOR CONTRIBUTIONS

Conceptualization, K.M.A., and J.L.G.; Methodology, K.M.A and J.L.G.; Investigation, K.M.A,W.S, K.M.K, C.A.F; Writing, K.M.A., and J.L.G.; Visualization, K.M.A.; Funding Acquisition, J.L.G.; Resources, J.L.G.; Supervision, K.M.A. and J.L.G.

## Materials and Methods

### Animal Maintenance

*C. elegans* were cultured using standard techniques. Upon thaw, worm strains were maintained un-starved at 20⁰ C for at least 3 - 6 generations prior to use. Each strain was maintained every 3-4 days by picking 4-5 hermaphrodites during their last larval stage, crescent L4(CL4). Males were initially propagated by isolating males from crowded or recently starved hermaphrodite plate and pairing them with a young unmated hermaphrodite. Thereafter, male plates were maintained in groups at a 3:1 ratio and maintained every 3-4 days. Animals were maintained on 6 cm plates seeded with 200 ml of OP50 grown overnight with agitation at 37⁰ C. Experimental animals were synchronized by either timed egg lay (1-2 hours) or picking the last larval stage (CL4). Unless otherwise indicated, synchronized animals were aged in groups of 50 and transferred every other day to avoid over-crowding and starvation. Assays were performed during peak reproduction (adult day 2) or reproductive decline (adult day 4). This study considered 96 hours post timed egg lay or 48 hours after the last larval stage (CL4) to be peak reproduction. Reproductive decline occurred 48 hours after peak reproduction.

### Fertility Assays

Assay Set-Up:

*Plate Preparation:*

3cm NGM plates were seeded with ∼30-50μl of OP50 (OD = 0.8) in the center of the plate 12-24 hours before use. Food was left to dry undisturbed to prevent OP50 from spreading to the walls of the plate. Lawns that are large or in contact with the edge of the plate decrease counting accuracy due to obscured visibility and experimenter fatigue. Plates were seeded the night before and stored upside down in a box at 4C (deli case) to prevent lawn thickening.

Thick lawns contribute to variable/inconsistent detection of unfertilized oocytes. Lawns that are too wet or thin increase animal leaving (crawling off the plate) or egg-retention. Plates were brought to room temperature 6-12 hours before use. All experimental plates were blinded at the start of the assay with GraphPad (https://www.graphpad.com/quickcalcs/randomize1).

*Male Propagation:*

Wild-type male plates were expanded by mating young adult males to young hermaphrodites at a 3:1 ratio approximately 72-96 hours before the start of the assay. In some cases, male maintenance plates were chunked to reduce density and risk of starvation.

#### Plate Set-Up: Early Reproduction (Adult Day 1-4)

Each hermaphrodite was singly housed and placed with 3 young adult males at crescent L4 following a 1–2-hour timed egg lay or picking directly from a maintenance plate. Mated hermaphrodites were transferred daily until death. Experimental plates were stacked in groups of 1-4, placed in metal trays and maintained at 20⁰ C. Unmated self-reproductive assays were set up the same way excluding the addition of males.

#### Plate Set-Up: Reproductive Decline (Adult Day 5-6)

Each hermaphrodite was singly housed and placed with 3 young adult males at the start of adult day 5 for 24 hours. Adult day 5 began 96 hours after CL4 or 144 hours following a 1–2-hour timed egg lay.

Mated hermaphrodites were transferred daily until death. Experimental plates were stacked in groups of 1-4, placed in metal trays and maintained at 20⁰ C. * Three of the nice experimental replicates were singled at crescent L4, and early unmated reproductive counts were collected prior to entering the mated assay at adult day 4-5.*

*Fertility Counts:*

After the mated hermaphrodites were transferred, plates were left at 20⁰ C or room temperature for an additional 12-24 hours before scoring unhatched and unfertilized eggs. Progeny were counted and scored for sex 48-72 hours after transfer. Progeny that failed to reach the last larval stage within 72 hours were not included. If progeny counts weren’t collected within 72 hours post-transfer, they were stored in the 4⁰ C for up to two additional days.

*Censorship and Matricide:*

Animals were censored on the day of incidence if they were severely damaged upon transfer, exposed to contamination, burrowed, or crawled onto the wall of the plate. No counts were collected on the day of censor, as they would underestimate the daily counts and bias the population. When NGM cracked or pulled away from the wall of the plate, care was taken to not mis-represent the counts. If too many animals were under the agar, the plate was censored on the day of incidence. Animals that died or had internal hatching were included until the day of occurrence. Death or censor type were recorded (non-responsive/dead or matricide). Counts collected on the day of death were not included in these analyses. Mating efficiency was determined for each hermaphrodite by calculating the proportion of all progeny that were male. Each mated animal had to generate at least 39% male by the end of the assay period to ensure adequate sperm transfer^212,213^. The frequency of male progeny in unmated hermaphrodites is less than 1%^214^. Hermaphrodites were considered incompletely mated if this criterion was not met. Hermaphrodites that were incompletely mated, censored, or died within 48 hours of mating were not included in the analysis. The data shown includes complete observations only.

### Reproductive stage bins

Reproductive Onset (L4 - D1): first 24 hours of adulthood, beginning at the last larval stage (CL4). Peak Reproduction (D1 - D3): two consecutive days following reproductive onset when egg-laying is maximal. The number of laid counts between unmated and mated animals are roughly equal.

Early Reproduction (L4-D4): First 4 days of the reproductive span.

Reproductive Decline (D5-D6): Two consecutive days following early reproduction when laid counts between unmated and mated differ most.

Late Reproduction (D7-D8): Two consecutive days following the reproductive decline when laid counts between unmated and mated are often equal.

*Metrics:*

Mating efficiency: The proportion of mated progeny that are male is at least 39%.

Viable progeny: Total number of mated progeny that reach the last larval stage within 72 hours of being laid per animal.

<u>Unfertilized eggs:</u> Total number of unfertilized eggs laid per animal. These eggs lack a definitive eggshell.

<u>Unhatched eggs:</u> Total number of fertilized eggs that do not hatch within 12-24 hours of being laid per animal. Those eggs have a definitive eggshell.

<u>Percent fertilized:</u> calculated by dividing the total number of fertilized counts (progeny and unhatched eggs) by the total number of laid counts (progeny, unfertilized eggs, unhatched eggs) per animal x 100. <u>Percent viable:</u> calculated by dividing the total number of progeny by the total number of fertilized counts (progeny, unhatched eggs) per animal x 100.

*Outlier Detection:*

Each replicate was normalized to the wild-type control and visualized in a violin plot using ggplot2 software in R Outliers were detected using the interquartile range set to moderate on pooled relative counts per variable. After outliers were removed, the remaining raw data was used to re-normalize to the wild-type control.

### Fluorescence Microscopy

*Live-Animal Mounting:*

Hermaphrodites were mounted with an eyelash or pick and paralyzed with 1-2μl of 0.167 mM levamisole on 15-well or 10-well multitest slide (VWR cat. no. IC096041505). The volume used was adjusted based on age:1μl was used for adult day 2 and 1.5 - 2μl for adult day 4. Minimal food was used to transfer the worm to limit debris detection during analysis.

*Image Acquisition:*

Images were acquired with Zeiss Axio Imager M2 microscope with a Zeiss Axiocam 503 monochrome camera at 10x and 20x magnification and exported in Tiff format. Exposure and power settings were optimized per experimental replicate. Generally, FAT-7::GFP exposure increased with age and VIT-2::GFP exposure decreased with age.

*Note for embryo quantification:* Two of the four early embryo datasets during peak reproduction were mounted in 1-2μl of 20% bleach for five minutes or less using the same multitest slides (VWR cat. no. IC096041505). to liberate eggs from the uterus. This protocol was used to ensure that in utero egg compaction did not explain the observed increase in yolk content.

#### FAT-7 Quantification

*FAT-7 Mated-Assay:*

Synchronized hermaphrodites were singled onto 3 cm mating plates during their last larval stage (CL4). Three young adult wild-type males were added to the plate and left for 24 hours. Mated hermaphrodites were removed from the mating plates and group housed (20-30 worms) on 6cm plates seeded 200μl of OP50 until adult day 2 (24 hours later). Unmated hermaphrodite controls were housed in groups of 30-50 on 6cm plates seeded with 300μl OP50 for the duration of the assay (CL4-adult day 2). An autofluorescence control was used during optimization to ensure that the exposure used wasn’t picking up artifacts.

*Image Analysis:*

Tiff images were analyzed using CellProfiler software. The intestinal ROI was determined by merging the DAPI and GFP channels and manually correcting the bounds of the intestine using the brightfield image as a guide. Total intestinal abundance of FAT-7 was determined by taking the integrated density of the intestine and dividing it by its area which is defined as the mean intensity of the object. Individual intestines were segmented into ten transverse regions using the WormToolbox module, StraightenWorms^215^. Each straightened intestine was saved in an output file to determine the orientation of the intestine and accuracy of the straightened intestine. Images that had debris or abnormal stretching were excluded from the region analysis. The un-straightened intestines were retained in the total intestinal measurements if no debris overlapped with the ROI. Periodically worms would burst during image acquisition, burst animals were not included in either dataset.

*Data Analysis:*

The data was exported from CellProfiler and aggregated into a single file. R was used to aggregate, perform calculations, visualize data, identify outliers and perform statistical testing. All measurements were reported on a per-animal basis. Statistical tests were performed on all pooled replicates after outliers were removed. Anatomical regions of the intestine were aggregated into three compartments to simplify comparison and interpretation. Each anatomical region represents the mean of those segments: anterior region (1-3), intermediate region (4-7), and posterior region (8-10).

*Outlier Detection:*

Figure 2B/3B/3E: Each replicate was normalized to the unmated wild-type control and visualized in a violin plot using ggplot2 software in R. Outliers were detected using a MAD Hampel filter on pooled relative FAT-7 intestinal abundance. Each replicate was normalized to the unmated wild-type control, and visualized in a violin plot using ggplot2 software in R. After outliers were removed, the remaining raw data was used to re-normalize to the unmated wild-type control.

Figure 2D-F and Figure 3C/F: Each replicate was normalized to the unmated wild-type control per region and visualized in a violin plot using ggplot2 software in R. Outliers were detected using the interquartile range set to moderate on pooled relative regional FAT-7 abundance. After outliers were removed, the remaining raw data was used to re-normalize to the unmated wild-type control.

#### VIT-2 Quantification

*Image Analysis:*

Tiff images were analyzed using CellProfiler. Intestinal, oocyte and egg ROIs were manually drawn using a merged brightfield and GFP guide. The mean was determined per object by dividing the total integrated intensity by the area of the object. Oocytes were categorized based on their position relative to the spermatheca/oviduct in the germline. The three most proximal oocytes were measured per arm when visible. The mean intensity of early embryos (< 5 cell divisions) and eggs (1-cell) were measured in the uterus. Periodically worms would burst during image acquisition, burst animals were not included in the intestinal or proximal oocyte datasets. Fertilized eggs were unaffected by bursting and were included. The mean or median of each oocyte position and egg measurement was reported on a per animal basis to avoid pseudo-replication.

*Data Analysis:*

The data was exported from CellProfiler and aggregated into a single file. R was used to aggregate, perform calculations, visualize data, identify outliers and perform statistical testing. All measurements were reported on a per-animal basis. Statistical tests were performed on all pooled replicates after outliers were removed.

*Outlier Detection:*

Each replicate was normalized to the unmated wild-type control and visualized in a violin plot using ggplot2 software in R. Outliers were detected using the interquartile range set to moderate on pooled relative VIT-2 abundance per tissue. After outliers were removed, the remaining raw data was used to re-normalize to the unmated wild-type control.

#### Progenitor Zone Staining and Quantification

Worms were synchronized with a 1–2-hour timed egg-lay. 48-56 hours later, 50-100 animals were transferred to a 6cm plate and aged to peak reproduction (48 hours post transfer) and reproductive decline (96 hours post transfer). On the day of staining, animals were washed off the plate with M9 into a 1.5ml Eppendorf tube and gravity settled. Supernatant was removed and the M9 was replaced. This process was repeated 3x. On the last wash the M9 was removed and replaced with 1ml of 0.2mM levamisole in PBS. Worms were transferred into a glass dish using a glass pipet. Worms were dissected by slicing behind the pharynx and the tail with a 0.5ml BD U-100 insulin syringes for up to 5 min. The dissection solution was removed from the glass dish and 2ml of fixative solution (3% PFA in 0.1M K2HPO4) was added directly to the dish. Gonads were fixed for 10 minutes. The fixative was removed and gonads were washed with 3ml of PBST (0.1% Tween-20) and transferred to a 5ml glass conical with a glass pipette. The conical was spun down at 1000xg for 30 seconds and the supernatant was removed. 2ml of ice cold 100% methanol was added, and the sample was incubated on ice for 5 minutes or stored in the −20⁰ C before staining. On the day of staining the sample was washed 3x with PBST (0.1% Tween-20), before incubating in 2ml of 100ng/ml DAPI in PBS for 5 minutes in the dark.

The sample was washed 2x with PBST (0.1% Tween-20) and mounted onto a glass slide with a 2% agarose pad. Fluoromount was added the coverslip and placed on the agarose pad. The slide was left to settle at room temperature in the dark for at least 4 hours or overnight before sealing with nail polish. Imaging took place within 1 day up to 2 weeks after preparation. Gonads were imaged at 20x with a 503 monochrome camera and a Zeiss Axio.Imager M2 microscope. The images were blinded using QuPath and the progenitor zone rows were counted by scrolling to the center of the gonad where the highest number of nuclei were visible. The number of rows were counted from the DTC to the transition zone, where at least two adjacent crescent moon shaped nuclei were visible.

Data Analysis:

The data was exported from QuPath and aggregated into a single file. R was used to aggregate, perform calculations, visualize data, identify outliers and perform statistical testing. All measurements were reported on a per-animal basis. Statistical tests were performed on all pooled replicates after outliers were removed.

*Outlier Detection:*

Each replicate was normalized to the wild-type control and visualized in a violin plot using ggplot2 software in R. Outliers were detected using the interquartile range set to moderate on pooled normalized samples. After outliers were removed, the remaining raw data was used to re-normalize to the median wild-type control.

### Nematocin signaling gene expression aging series

#### Worm culture

Wild-type(N2) worms were grown and aged on 10cm plates and incubated at 20°C. At each timepoint, worms were washed off the plate with M9, gravity settled and washed 3x to remove progeny and bacteria. The supernatant was removed on the last wash and 500ml Trizol was added before the sample was flash frozen with liquid nitrogen and stored in the −80⁰ C for several months.

#### RNA Isolation

Prior to RNA extraction, the sample was lysed by undergoing 6-7 freeze thaw cycles rotating between liquid nitrogen, 37⁰ C water-bath and 30 second vortex. These steps were repeated until worm corpses were mostly dissolved (7 cycles). RNA was isolated with the Zymo Direct-zol RNA Miniprep Kit (including on-column DNase treatment), and RNA concentration and purity were assessed with a Nanodrop ND-1000 spectrophotometer. If low RNA purity was detected, samples were cleaned using the Qiagen RNeasy Plus Micro Kit. N2 worms had an average RNA concentration of 180.8 ng/uL, (standard deviation: 158.0 ng/uL) and therefore an average total RNA yield of 9.04 ug (standard deviation: 7.902 ug). A260/280 ratios for N2 samples averaged 2.10 with a standard deviation of 0.05. RNA integrity was assessed by running a representative subsample through the Agilent Bioanalyzer 2100 RNA 6000 Nano Kit. RIN numbers averaged 9.9, with a standard deviation of .13.

#### cDNA Synthesis

cDNA was generated using the RT-qPCR protocol for Thermo Scientific Maxima H Minus First Strand cDNA Synthesis Kit, with dsDNase. ≤1 ug of RNA was used in each well, with wells brought to exactly 1 ug RNA if possible. Total reaction volume was 20 uL. Reaction was primed with Oligo(dT)18 primers and random hexamer primers, each at 100uM. Reaction was assembled on ice, except for 2 min at 37°C for gDNA elimination, and 5 min at 65°C for secondary RNA structure elimination. Then cDNA was generated by incubations of 10 min at 25°C, 15 min at 50°C, and 5 min at 85°C. No Reverse Transcriptase (NRT) controls were performed and checked later by qPCR. No NRT control samples showed amplification qPCR. cDNA samples were stored at −80°C.

#### qPCR

qPCR was performed using the Bio-Rad iTaq Universal Probes Supermix kit, containing antibody-mediated hot-start iTaq DNA polymerase, and using 10 uL reaction volume with .1 uL of cDNA per well. Each age point was analyzed using three sets of biological replicates, and each biological replicate was plated as a set of three technical replicates. Reaction was set up manually in clear well plates, using Bio-Rad Clear 96 Well Plates model HSP 9601 or Thermo Scientific AB-1384 clear 384 well plates. qPCR was run on a Bio-Rad CFX96 Touch or CFX 384 Touch thermocycler. Thermocycling parameters were 1: 95°C for 30s, 2: 95°C for 3s, 3: 60°C for 30s, then repeat steps 2 & 3 40x. No Template controls were included as a negative control for contamination. All qPCR assays used were commercially produced probe-based Thermo Fisher TaqMan gene expression assays, using FAM flourophores and minor groove binder nonflourescent quenchers. Forward and reverse primers were at a concentration of 900nM each, and the probe was at 250nM.

#### Statistical analysis

Relative gene expression was analyzed using Bio-Rad CFX Maestro for Mac 1.1, Version 4.1.2434.0124. Cq values were determined using the Regression method. Technical replicates were subjected to Grubb’s Test for Outliers to eliminate individual wells as necessary. Technical Replicate groups were excluded and repeated if the standard deviation of the three Cq values exceeded .5.

Relative gene expression was assessed using the ΔΔCq method, normalized to 4 reference genes. Act-2, Ama-1, Cdc-42, and Pmp-3 were selected as reference genes, based on a literature search.

Reference gene stability was confirmed using the CFX Maestro Reference Gene Selection tool. Statistical significance was based on a p-value of ≤.05.

### Single-cell Expression Dataset

Data from Roux et al., 2023 was analyzed in R using Seurat v5. Dotplots for gene expression by cell type were generated using day 1-15.

**Table 1:**
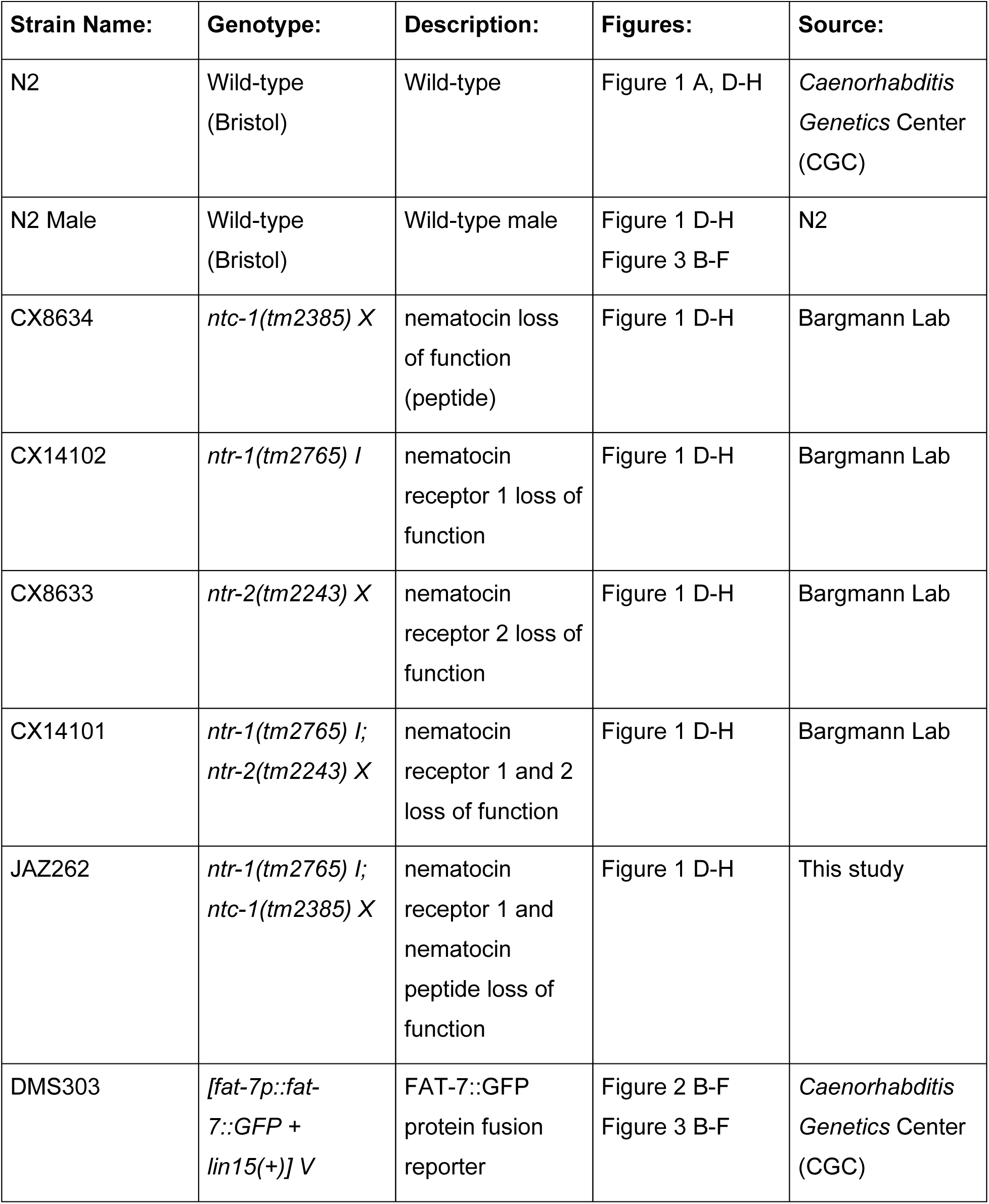
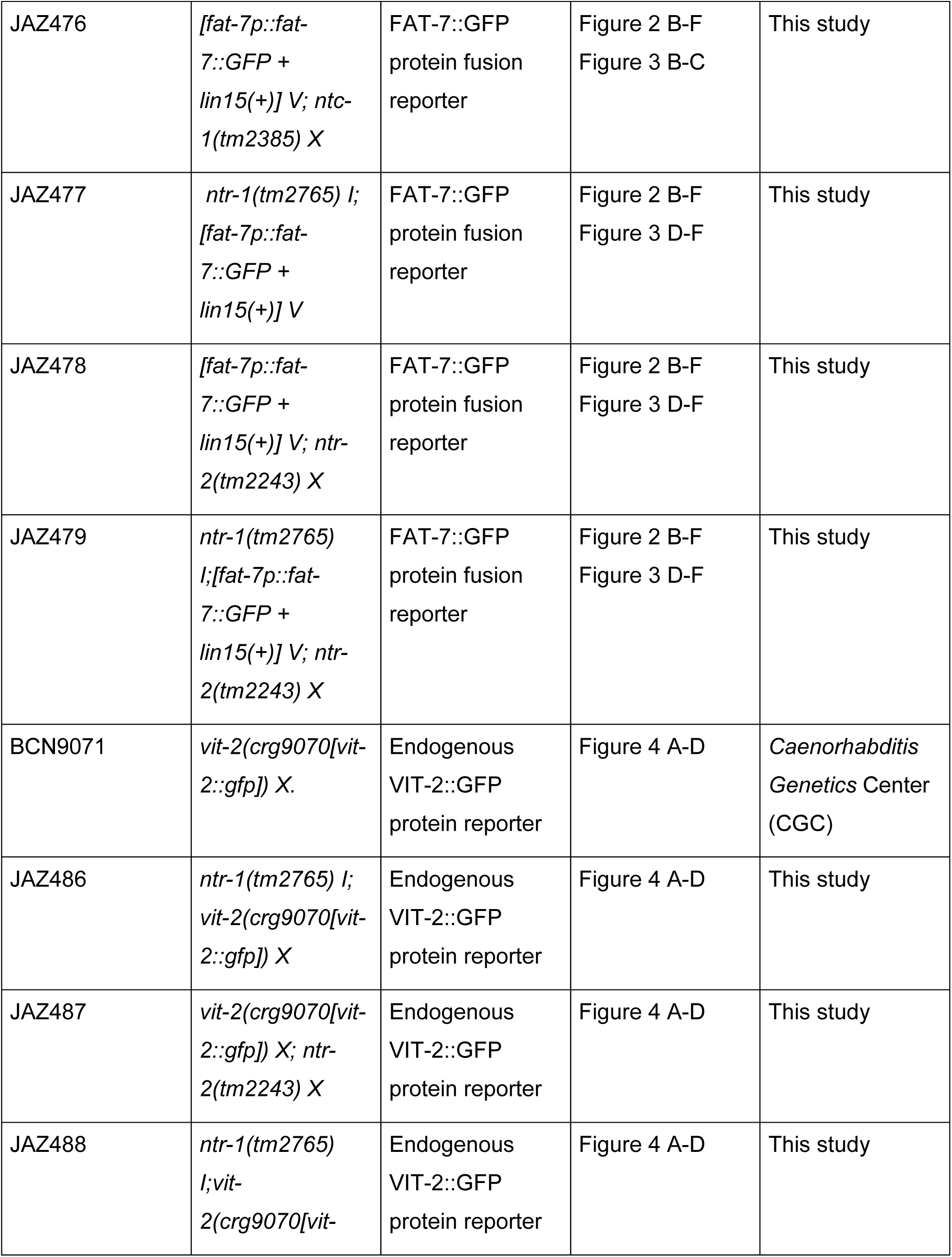
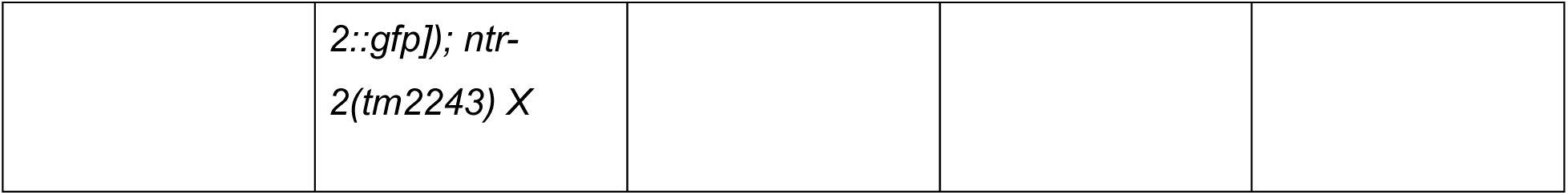
Experimental Models.

**Table 2:**
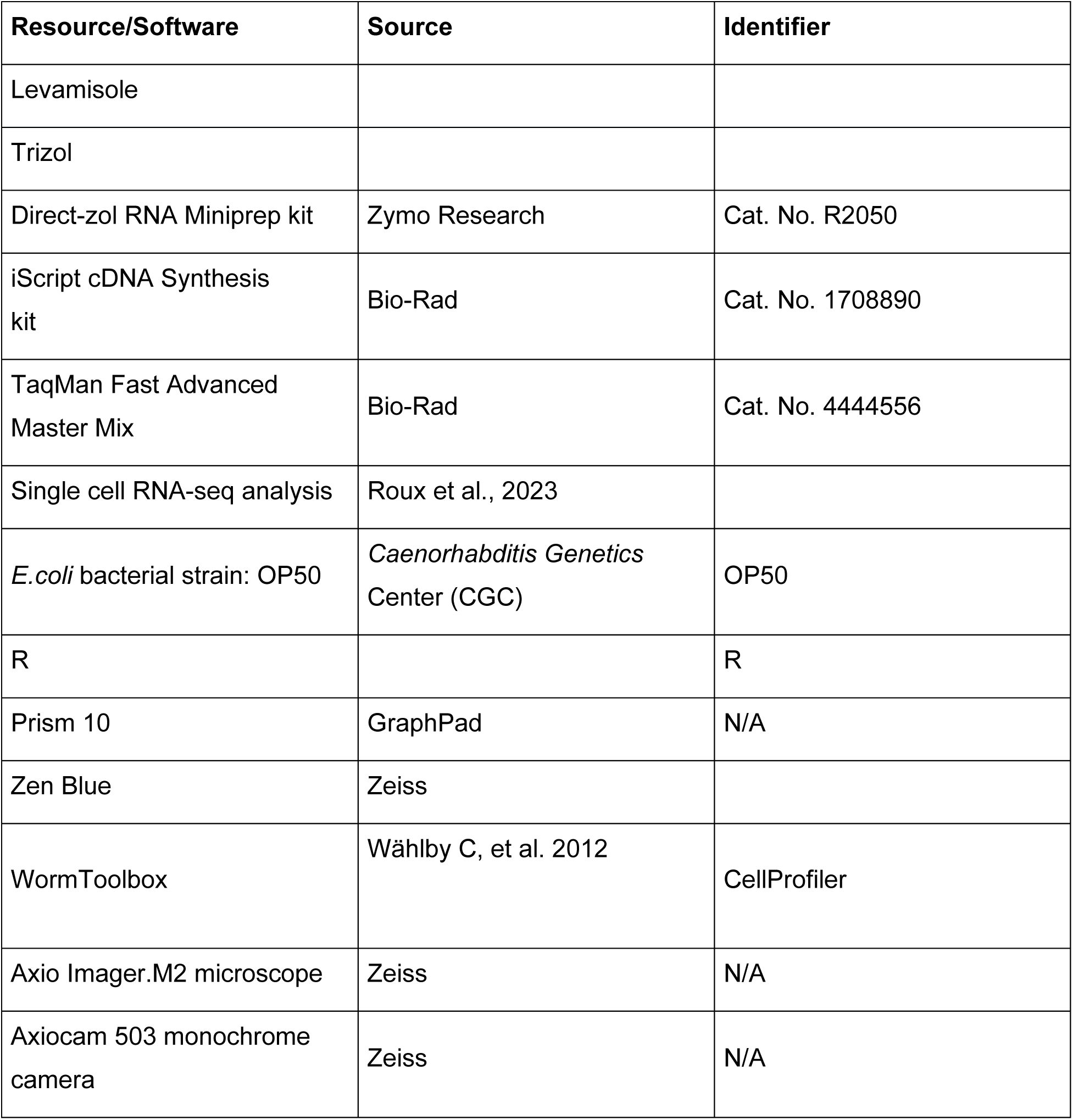
Resources.

**Table 3:**
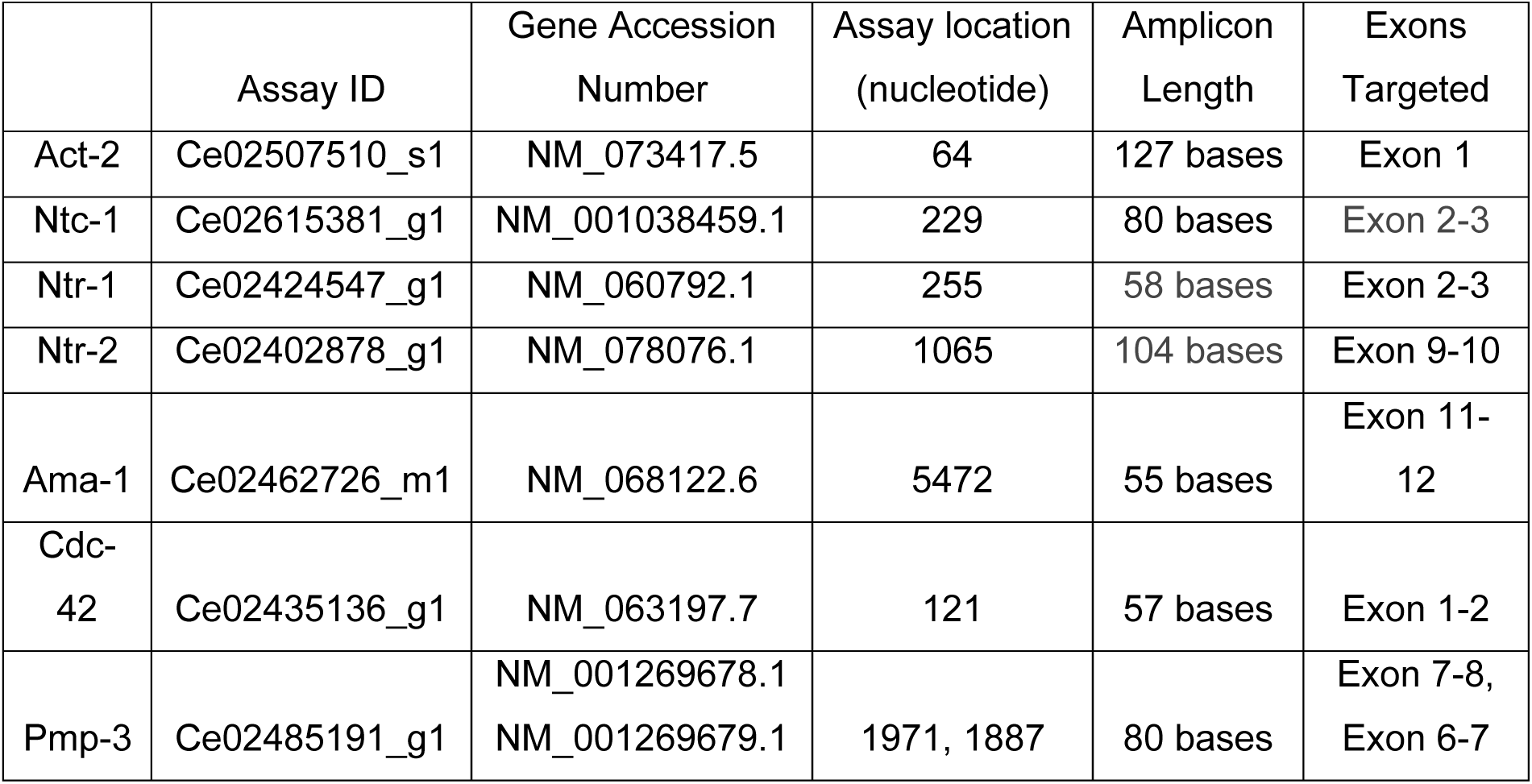
qPCR primers.

---

**Supplemental Figure 1:**
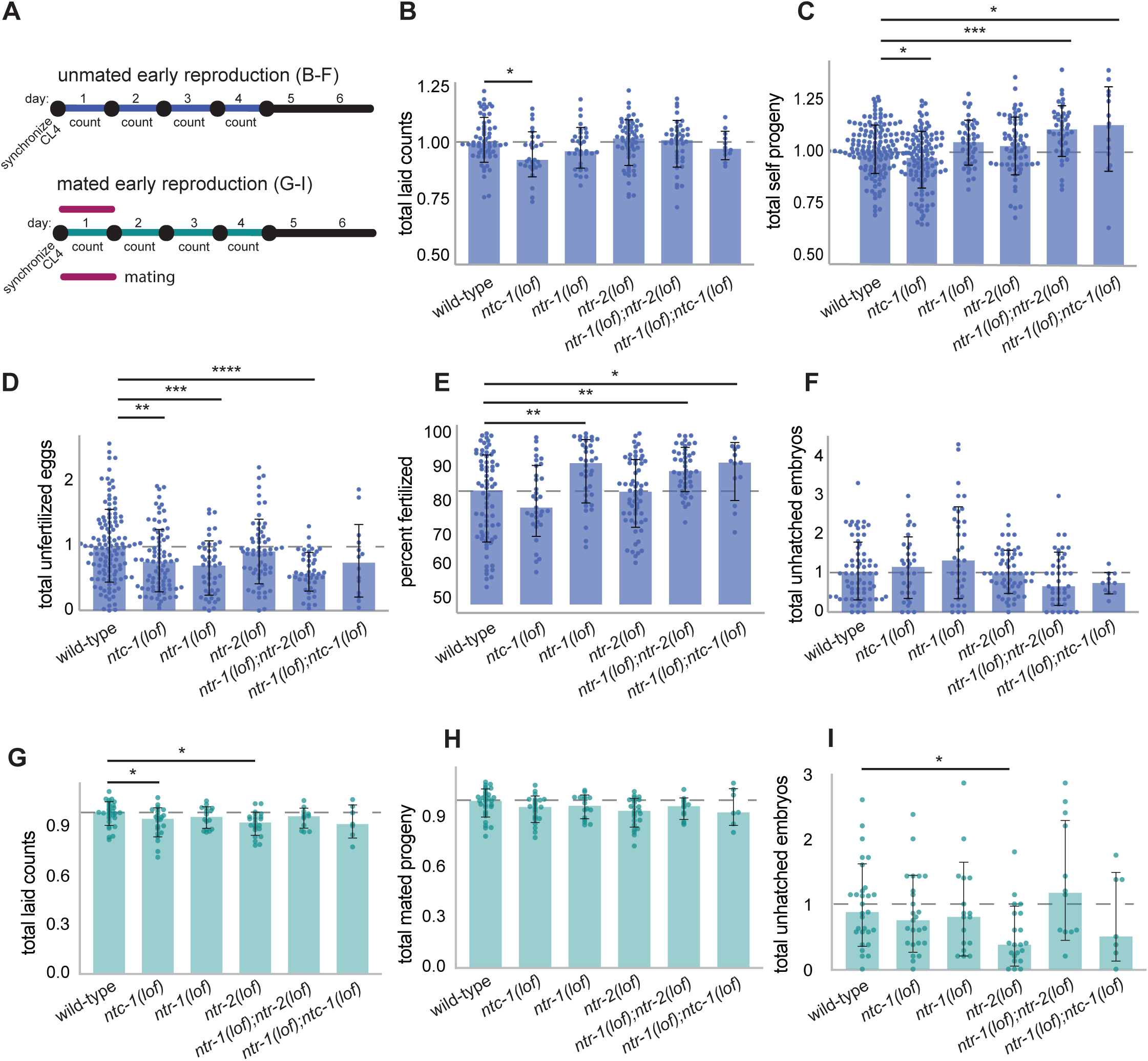
**(A)** Experimental timeline for unmated and mated early reproduction with figure designation. **(B) Unmated Early Reproduction (1- 4):** Relative number of laid counts (self-progeny, unfertilized eggs and unhatched eggs) during the first 4 days of adulthood. Kruskal- Wallis + Dunn Test + Benjamini-Hochberg (FDR) corrections were applied to pooled samples between mutant genotype compared to wild-type only. All significant comparisons and adjusted p-values are shown. Adjusted *p*-values indicated by * *p* < 0.05, ** *p* <0.01, *** *p* < 0.001, **** *p* < 0.0001. Number of experimental and biological replicates: (N = 6, 3, 3, 4, 3, 2; n = 61, 29, 33, 60, 41, 12) **(C) Unmated Early Reproduction (1- 4):** Relative number of self-progeny during the first 4 days of adulthood. Kruskal- Wallis + Dunn Test + Benjamini-Hochberg (FDR) corrections were applied to pooled samples between mutant genotype compared to wild-type only. All significant comparisons and adjusted p-values are shown. Adjusted *p*-values indicated by * *p* < 0.05, ** *p* <0.01, *** *p* < 0.001, **** *p* < 0.0001. Number of experimental and biological replicates: (N = 9, 6, 3,4,3, 2; n = 141, 110, 43, 66, 43, 14) **(D) Unmated Early Reproduction (1- 4):** Relative number of unfertilized eggs during the first 4 days of adulthood. Kruskal- Wallis + Dunn Test + Benjamini-Hochberg (FDR) corrections were applied to pooled samples between mutant genotype compared to wild-type only. All significant comparisons and adjusted p-values are shown. Adjusted *p*-values indicated by * *p* < 0.05, ** *p* <0.01, *** *p* < 0.001, **** *p* < 0.0001. Number of experimental and biological replicates: (N = 8, 5, 3, 4, 3, 2; n = 119, 83, 46, 62, 44, 14) **(E) Unmated Early Reproduction (1- 4):** Percent of laid counts that were fertilized during the first 4 days of adulthood. Kruskal- Wallis + Dunn Test + Benjamini-Hochberg (FDR) corrections were applied to pooled samples between mutant genotype compared to wild-type only. All significant comparisons and adjusted p-values are shown. Adjusted *p*-values indicated by * *p* < 0.05, ** *p* <0.01, *** *p* < 0.001, **** *p* < 0.0001. Number of experimental and biological replicates: (N = 6, 6, 3,4,3, 2; n = 72, 33, 36, 61, 44, 13) **(F) Unmated Early Reproduction (1- 4):** Relative number of unhatched fertilized eggs during the first 4 days of adulthood. Kruskal- Wallis + Dunn Test + Benjamini-Hochberg (FDR) corrections were applied to pooled samples between mutant genotype compared to wild-type only. All significant comparisons and adjusted p-values are shown. Adjusted *p*-values indicated by * *p* < 0.05, ** *p* <0.01, *** *p* < 0.001, **** *p* < 0.0001. Number of experimental and biological replicates: (N = 6, 3, 3,4,3, 2; n = 69, 33, 36, 58, 41, 11) **(G) Young Mated Early Reproduction (1- 4):** Relative number of laid counts (mated progeny, unfertilized eggs and unhatched eggs) during the first 4 days of adulthood. Kruskal- Wallis + Dunn Test + Benjamini-Hochberg (FDR) corrections were applied to pooled samples between mutant genotype compared to wild-type only. All significant comparisons and adjusted p-values are shown. Adjusted *p*-values indicated by * *p* < 0.05, ** *p* <0.01, *** *p* < 0.001, **** *p* < 0.0001. Number of experimental and biological replicates: (N = 3, 3, 2, 3, 2, 1; n = 31, 24, 16, 23, 12, 7) **(H) Young Mated Early Reproduction (1 - 4):** Relative number of mated progeny during the first 4 days of adulthood. Kruskal- Wallis + Dunn Test + Benjamini-Hochberg (FDR) corrections were applied to pooled samples between mutant genotype compared to wild-type only. All significant comparisons and adjusted p-values are shown. Adjusted *p*-values indicated by * *p* < 0.05, ** *p* <0.01, *** *p* < 0.001, **** *p* < 0.0001. Number of experimental and biological replicates: N = 3, 3, 2, 3, 2, 1; n = 31, 23, 16, 24, 12, 7 **(I) Young Mated Early Reproduction (1 - 4):** Relative number of unhatched fertilized embryos during the first 4 days of adulthood. Kruskal- Wallis + Dunn Test + Benjamini-Hochberg (FDR) corrections were applied to pooled samples between mutant genotype compared to wild-type only. All significant comparisons and adjusted p-values are shown. Adjusted *p*-values indicated by * *p* < 0.05, ** *p* <0.01, *** *p* < 0.001, **** *p* < 0.0001. Number of experimental and biological replicates: N = 3, 3, 2, 3, 2, 1; n = 30, 25,1 7, 21, 12, 7

**Supplemental Figure 2:**
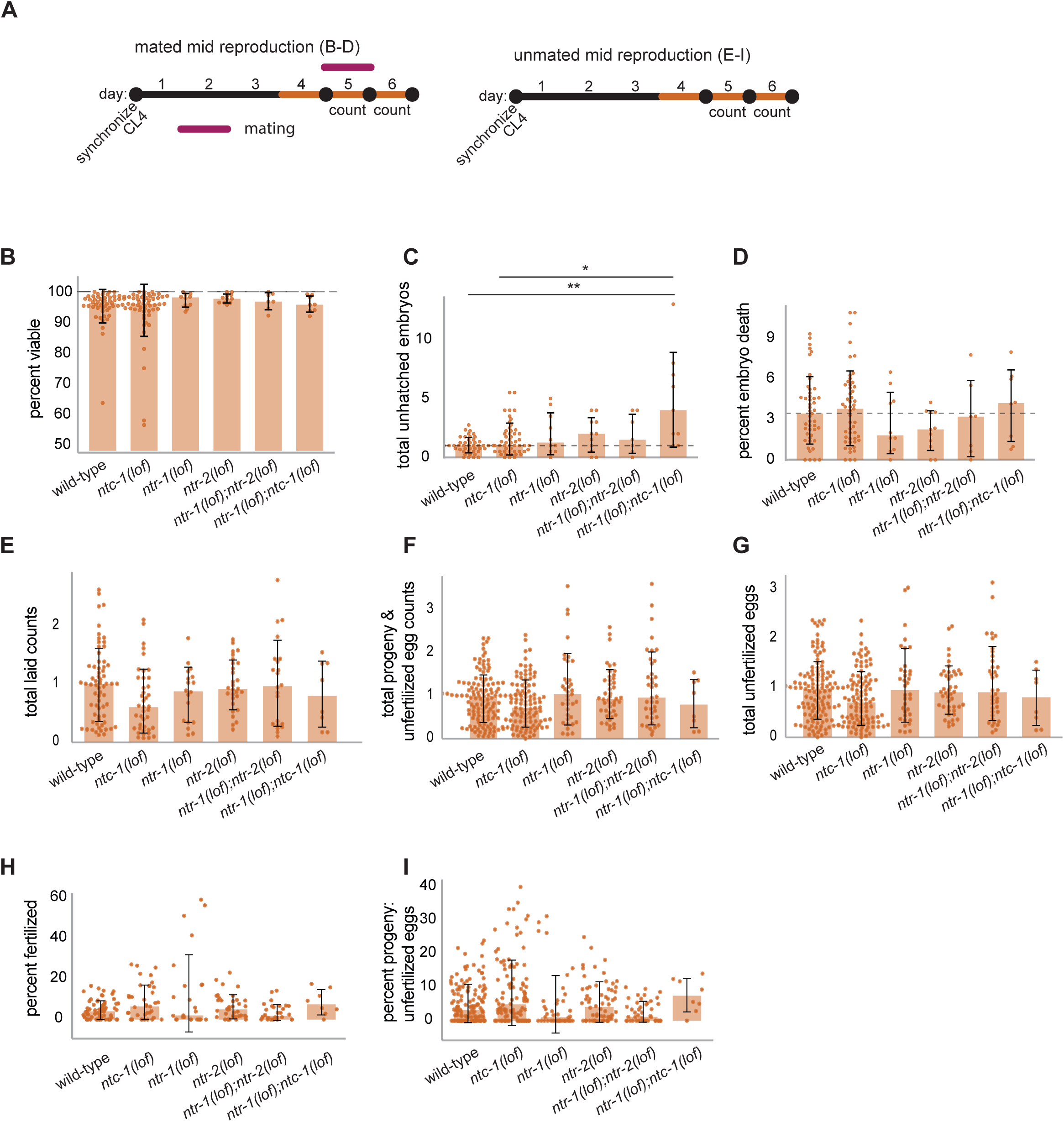
**(A)** Experimental timeline for unmated and mated conditions during the reproductive decline with figure designation. **(B) Mated D4-D5:** Percent of fertilized eggs that are viable from adult days 5-6. Kruskal- Wallis + Dunn Test + Benjamini-Hochberg (FDR) corrections were applied to pooled samples comparing all genotype interactions. All significant comparisons and adjusted p-values are shown. Adjusted *p*-values indicated by * *p* < 0.05, ** *p* <0.01, *** *p* < 0.001, **** *p* < 0.0001. Number of experimental and biological replicates: (N = 4, 4, 1, 2, 1, 1; n = 50, 55,10,11, 7, 8). *Note: no outlier removal* **(C) Mated D4-D5:** Relative total median mated unhatched embryos from adult days 5-6. Kruskal-Wallis + Dunn Test + Benjamini-Hochberg (FDR) corrections were applied to pooled samples comparing all genotype interactions. All significant comparisons and adjusted p-values are shown. Adjusted *p*-values indicated by * *p* < 0.05, ** *p* <0.01, *** *p* < 0.001, **** *p* < 0.0001. Number of experimental and biological replicates: (N= 4, 4, 1, 2, 1, 1; n = 45, 51, 10, 11, 7, 9) **(D) Mated D4-D5:** Percent of fertilized eggs that are dead from adult days 5-6. Kruskal- Wallis + Dunn Test + Benjamini-Hochberg (FDR) corrections were applied to pooled samples comparing all genotype interactions. All significant comparisons and adjusted p-values are shown. Adjusted *p*-values indicated by * *p* < 0.05, ** *p* <0.01, *** *p* < 0.001, **** *p* < 0.0001. Number of experimental and biological replicates: (N= 4, 4, 1, 2, 1, 1; n = 47, 51, 10, 11, 7, 8) **(E) Unmated Reproductive Decline (4 - 6):** Relative number of laid counts (self-progeny, unfertilized eggs and unhatched eggs) during adult days 5-6. All samples were normalized to the median wild-type control. Kruskal- Wallis + Dunn Test + Benjamini-Hochberg (FDR) corrections were applied to pooled samples comparing mutant genotype to wild-type only. All significant comparisons and adjusted p-values are shown. Adjusted *p*-values indicated by * *p* < 0.05, ** *p* <0.01, *** *p* < 0.001, **** *p* < 0.0001. Number of experimental and biological replicates: (N = 7, 5, 2, 3, 2, 1; n = 66, 41, 18, 29, 22, 8). **(F) Unmated Reproductive Decline (4 - 6):** Relative number of self-progeny and unfertilized eggs during adult days 5-6 Kruskal- Wallis + Dunn Test + Benjamini-Hochberg (FDR) corrections were applied to pooled samples comparing mutant genotype to wild-type only. All significant comparisons and adjusted p-values are shown. Adjusted *p*-values indicated by * *p* < 0.05, ** *p* <0.01, *** *p* < 0.001, **** *p* < 0.0001. Number of experimental and biological replicates: (N = 11, 9, 3, 4, 3, 1; n = 128, 104, 37, 45, 39, 8) **(G) Unmated Reproductive Decline (4 - 6):** Relative number of unfertilized eggs during adult days 5-6. Kruskal- Wallis + Dunn Test + Benjamini-Hochberg (FDR) corrections were applied to pooled samples between mutant genotype compared to wild-type only. All significant comparisons and adjusted p-values are shown. Adjusted *p*-values indicated by * *p* < 0.05, ** *p* <0.01, *** *p* < 0.001, **** *p* < 0.0001. Number of experimental and biological replicates: (N = 11, 9, 3, 4, 3, 1; n = 128, 104, 37, 45, 39, 8) **(H) Unmated Reproductive Decline (4 - 6):** Percent of laid counts that are fertilized during adult days 5-6. Kruskal- Wallis + Dunn Test + Benjamini-Hochberg (FDR) corrections were applied to pooled samples between mutant genotype compared to wild-type only. All significant comparisons and adjusted p-values are shown. Adjusted *p*-values indicated by * *p* < 0.05, ** *p* <0.01, *** *p* < 0.001, **** *p* < 0.0001. Number of experimental and biological replicates: (N = 7, 5, 2, 3, 2, 1; n = 67, 39, 25, 38, 30, 8) **(I) Unmated Reproductive Decline (4 - 6):** Percentage of progeny & unfertilized eggs that are progeny during adult days 5-6. Kruskal- Wallis + Dunn Test + Benjamini-Hochberg (FDR) corrections were applied to pooled samples between mutant genotype compared to wild-type only. All significant comparisons and adjusted p-values are shown. Adjusted *p*-values indicated by * *p* < 0.05, ** *p* <0.01, *** *p* < 0.001, **** *p* < 0.0001. Number of experimental and biological replicates: (N = 12, 10, 4, 6, 4, 1; n = 138, 108, 42, 64, 45, 8)

**Supplemental Figure 3:**
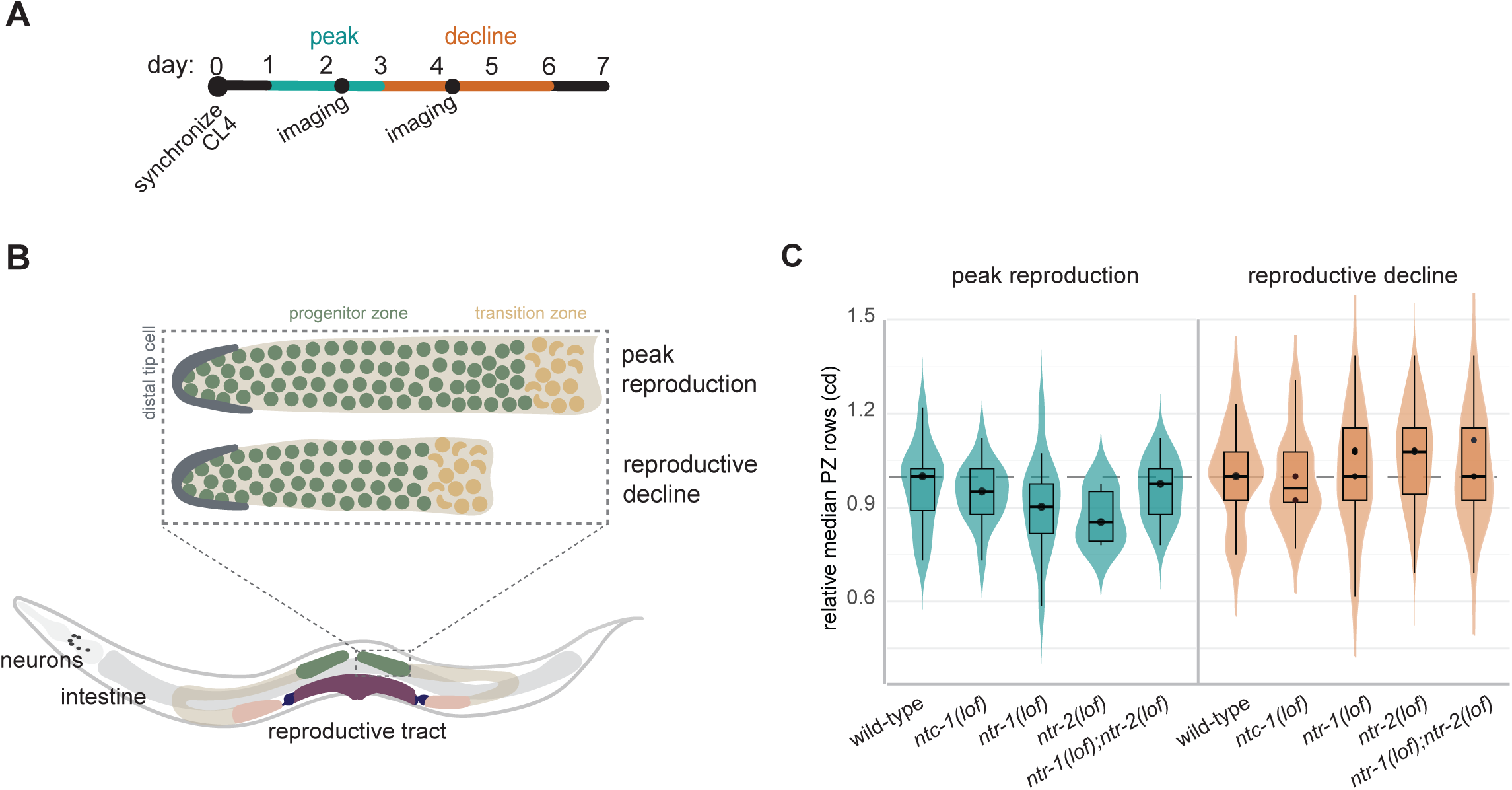
**(A) Experimental timeline**: germline dissection, staining and imaging of unmated worms during reproductive peak and decline with figure designation. **(B) Schematic of PZ:** Representative diagram of the distal gonad. The distal tip cell shown in dark green, progenitor cells shown in green and the transition zone shown in dark yellow. The number of green cell rows (progenitor zone) leading up to crescent moon shaped nuclei (dark yellow) are compared between genotypes and decline with age. During peak reproduction the mean number of rows is ∼ 20 and declines to ∼ 13 by the reproductive decline in the unmated condition. **(C) Unmated PZ:** Relative total unmated progenitor zone rows per animal during peak reproduction(D2-D3) and reproductive decline (D4-D5). All conditions were normalized to the median wild-type control. Kruskal- Wallis + Dunn Test + Benjamini-Hochberg (FDR) corrections were applied to pooled samples comparing each mutant genotype to wild-type only. All significant comparisons and adjusted p-values are shown. Adjusted *p*-values indicated by * *p* < 0.05, ** *p* <0.01, *** *p* < 0.001, **** *p* < 0.0001. Number of experimental and biological replicates: Peak reproduction: N = 1, 1,1,1,1; n = 14, 16, 16, 6, 20 and Reproductive decline: N = 3, 3, 3, 3, 2; n = 61, 66, 77, 89, 45.

**Supplemental Figure 4:**
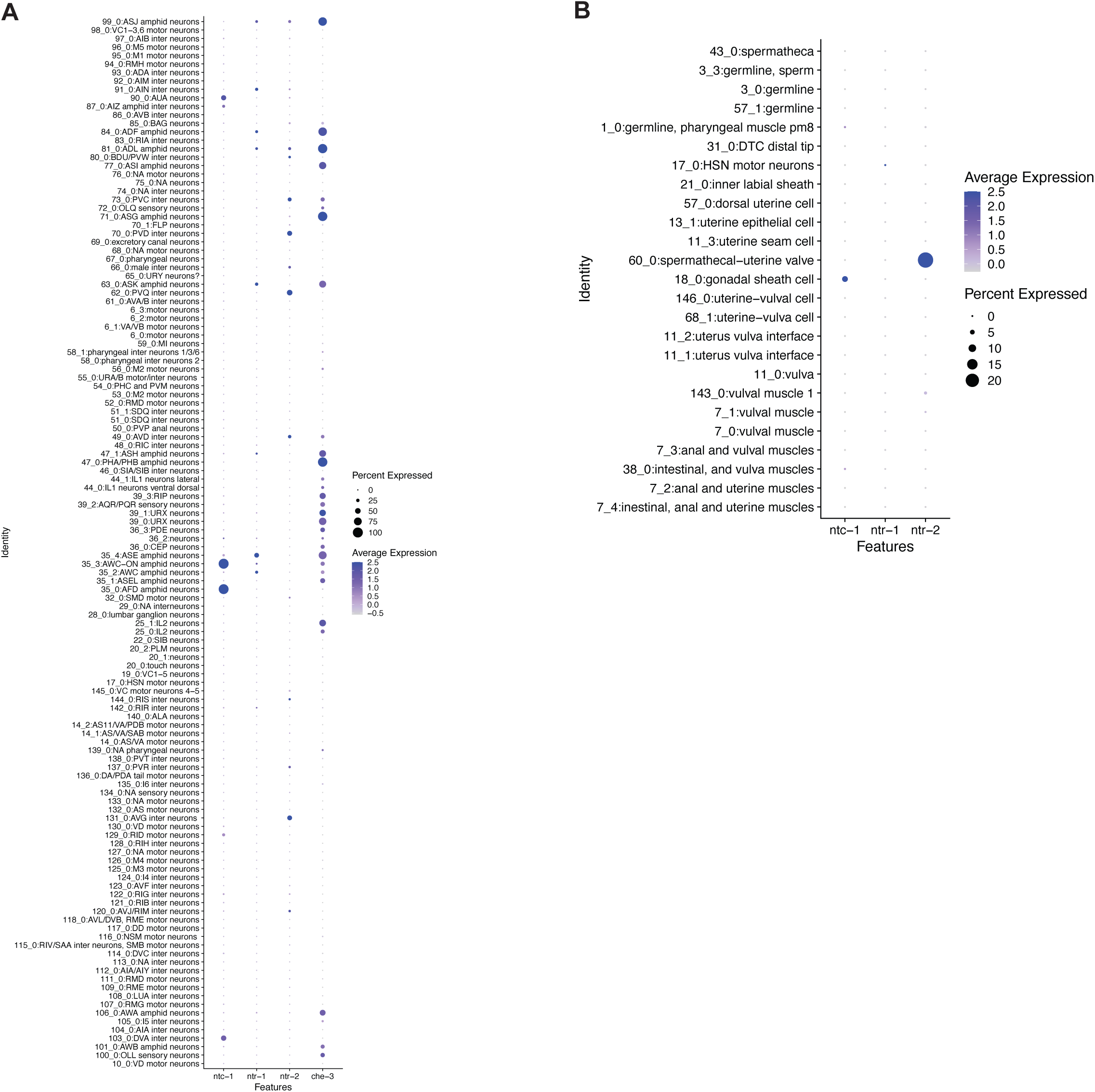
**(A)** Single cell dot plots of neurons normalized to the relative expression of *ntc-1*, *ntr-1*, *ntr-2,* and *che-3* across adult days 1-15. Data obtained from Calico aging sc-RNA-seq dataset.^1^ **(B) Single cell dot plots of reproductive tissues** normalized to the relative expression of *ntc-1*, *ntr-1*, and *ntr-2* across adult days 1-15. Data obtained from Calico aging sc-RNA-seq dataset. ^1^

**Supplemental Figure 5:**
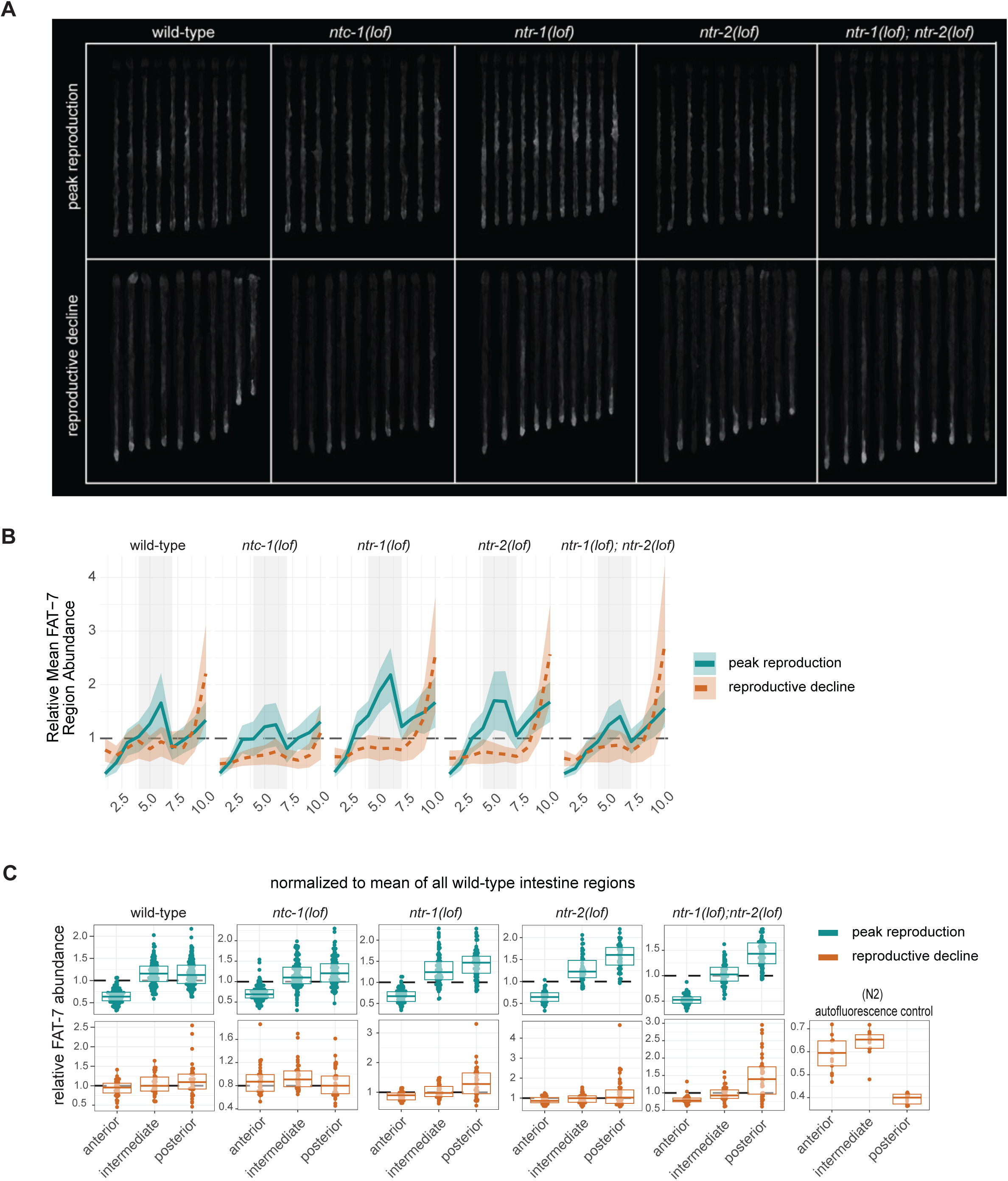
(A) Representative intestinal images of FAT-7 abundance in unmated animals. Images represent a single cohort. Data represented in plots below. **(B) Representative FAT-7 abundance and distribution relative to wildtype per age.** Relative abundance of FAT-7 across the segments of the intestine represented as a line plot where the mean (solid line) and standard error of the mean (ribbon) are shown per condition. Shaded grey background represents the intermediate intestine. Each condition was normalized to the respective wildtype intestine age. Data shown from single replicate. No statistical testing applied. **(C) Relative FAT-7 abundance by intestinal region (all replicates shown).** Aggregated segments are plotted adjacent to demonstrate regional bias across the intestine with age. All values are normalized to the mean aggregate of the unmated WT intestine. No statistical testing applied.

**Supplemental Figure 6:**
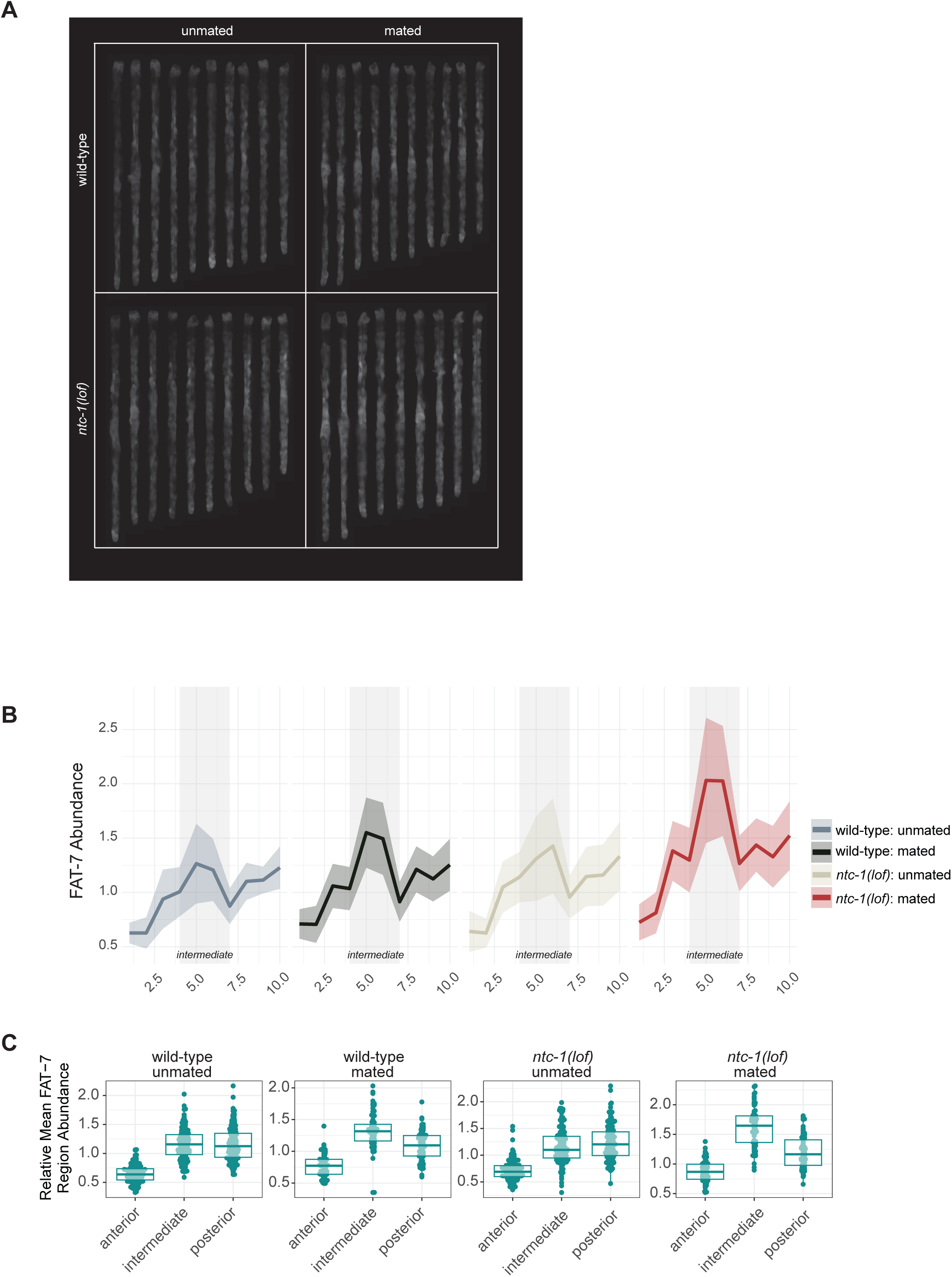
(A) Representative intestinal images of. unmated and mated wild-type and *ntc-1(lof)* animals from a single replicate. **(B) Representative FAT-7 abundance and distribution relative to unmated wildtype.** Relative abundance of FAT-7 across the segments of the intestine represented as a line plot where the mean (solid line) and standard error of the mean (ribbon) are shown per condition. Each condition was normalized to the unmated wildtype intestine. Data shown from single replicate. No statistical testing applied. **(D) Relative FAT-7 abundance by intestinal region.** Aggregated segments are plotted adjacent to demonstrate regional bias across the intestine with age. All values are normalized to the mean aggregate of the unmated WT intestine. No statistical testing applied.

**Supplemental Figure 7:**
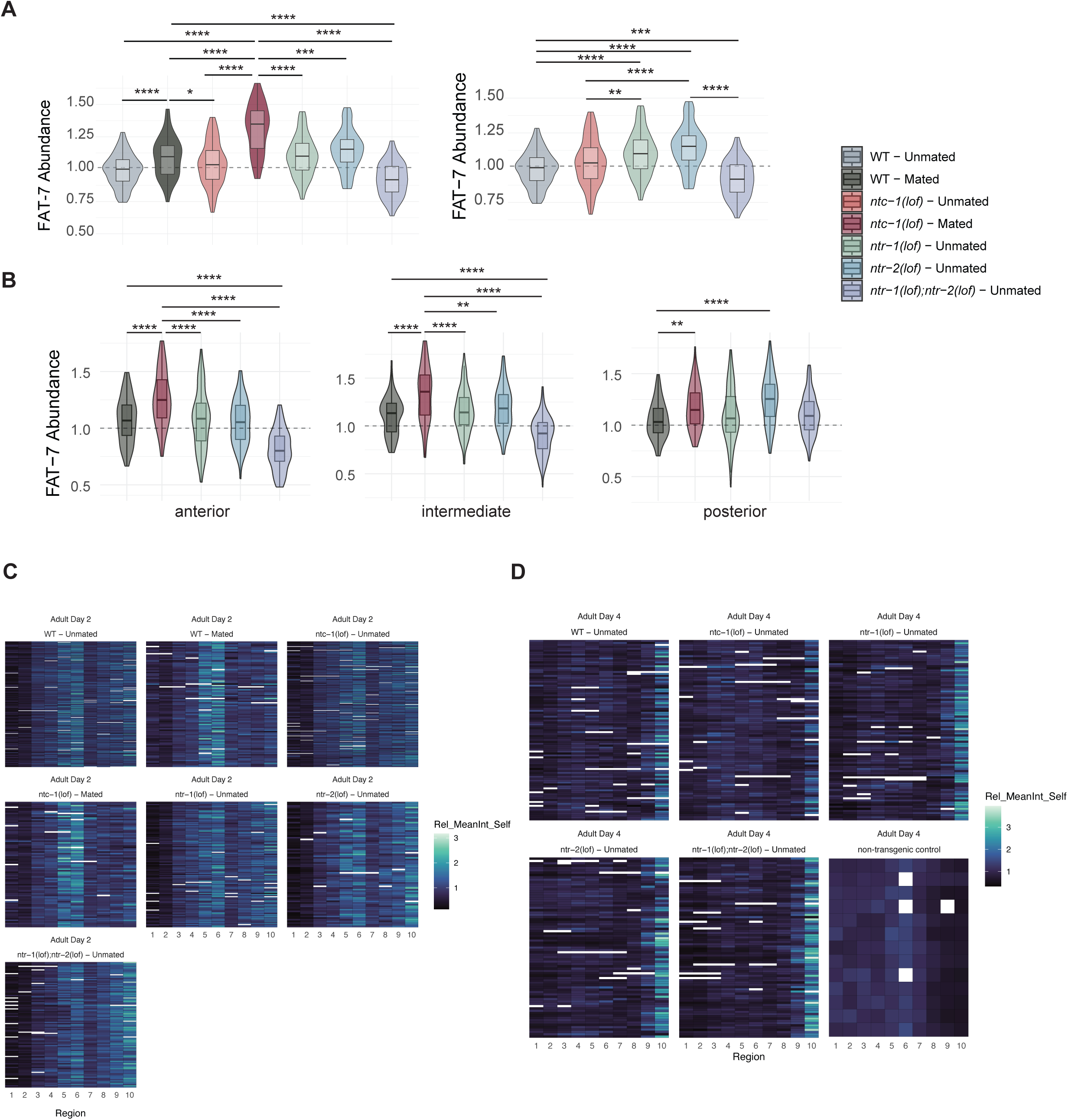
(A) Relative median abundance of intestinal FAT-7 during peak reproduction in unmated and mated animals. All animals were normalized to the median unmated wild-type control. Kruskal-Wallis + Dunn Test + Benjamini-Hochberg (FDR) corrections were applied to pooled samples between all conditions. All significant comparisons and adjusted p-values are shown. Adjusted *p*-values indicated by * *p* < 0.05, ** *p* <0.01, *** *p* < 0.001, **** *p* < 0.0001. Number of experimental and biological replicates: N = 9, 4, 9, 4, 4, 4, 3, 4; n = 192, 169, 197, 67, 106, 73, 105. Medians: 1.00, 1.10, 1.03, 1.35, 1.10, 1.15, 0.92. **(B) Relative mean abundance of FAT-7 per region** during peak reproduction in unmated and mated animals. All animals were normalized to the median unmated wild-type control. 1-way ANOVA + TukeyHSD were applied. All significant comparisons and adjusted p-values are shown. Adjusted *p*-values indicated by * *p* < 0.05, ** *p* <0.01, *** *p* < 0.001, **** *p* < 0.0001. Number of experimental and biological replicates: N = 9, 4, 9, 4, 4, 3, 4; n = 200-203, 71, 202-205, 67-68, 109-112, 74-75, 101-103. (C) **Heatmaps of FAT-7 distribution across adult day 2 intestines per condition.** The intensity was normalized to the mean of each worms abundance to show localization independent of relative abundance to wild-type. Intestine values are ordered chronologically from the anterior to posterior on the x-axis. White spaces are missing values due to outlier removal. (D) **Heatmaps of FAT-7 distribution across adult day 4 intestines per condition.** The intensity was normalized to the mean of each worms abundance to show localization independent of relative abundance to wild-type. Intestine values are ordered chronologically from the anterior to posterior on the x-axis. White spaces are missing values due to outlier removal.

